# Cellular context shapes the transcriptomic impact of somatic variants

**DOI:** 10.64898/2026.10.28.760887

**Authors:** Peisen Sun, Peng Jia, Chuyuan Lou, Xiaona Li, Miaojing Li, Ruifen Zhang, Hui Lin, Xiujuan Li, Yunsong Zhou, Shuzheng Cai, Zexin Zhang, Pengyu Zhang, Tun Xu, Songbo Wang, Yudong Wang, Mingxuan Li, Yuyan Gong, Guoqin Sang, Jie Yin, Jizhe Li, Qi Xie, Zongsheng Yue, Dan Xu, Xiaofei Yang, Stephen J. Bush, Songmei Geng, Deyu Meng, Xuexin Li, Kai Ye

**Author notes:** Correspondence (K.Y.) and (X.X.L.). These authors contributed to the work equally (Peisen Sun, Peng Jia, Chuyuan Lou).

## Abstract

Somatic SNVs are commonly prioritized by recurrence or sequence annotation, leaving the molecular consequences of private variants unresolved. We developed PrismSNV and Ligo-Seq, a single-cell full-length transcriptome sequencing platform, to quantify transcriptomic displacement associated with within-cell SNV-state toggling. Across 26 samples from three cancer types, more than 60% of analyzed SNVs showed significant cell-type-specific displacement with over 30% variation in magnitude. Sequence-based deleteriousness was poorly aligned with cellular displacement, and fewer than 3% of top-displacement SNVs across three tumor types were catalogued in ClinVar. Displacement was strongest in transcriptionally plastic, less differentiated states. Analysis of 655,349 CD4+ T-cell transcriptomes further showed that SNV-gene associations changed in magnitude and sometimes direction during activation. Lack of recurrence or clinical annotation does not imply transcriptomic inertness; variant impact is shaped by cellular context rather than sequence alone, establishing the variant-cell pair as a critical unit for interpreting somatic variation.

**Graphical Abstract:** 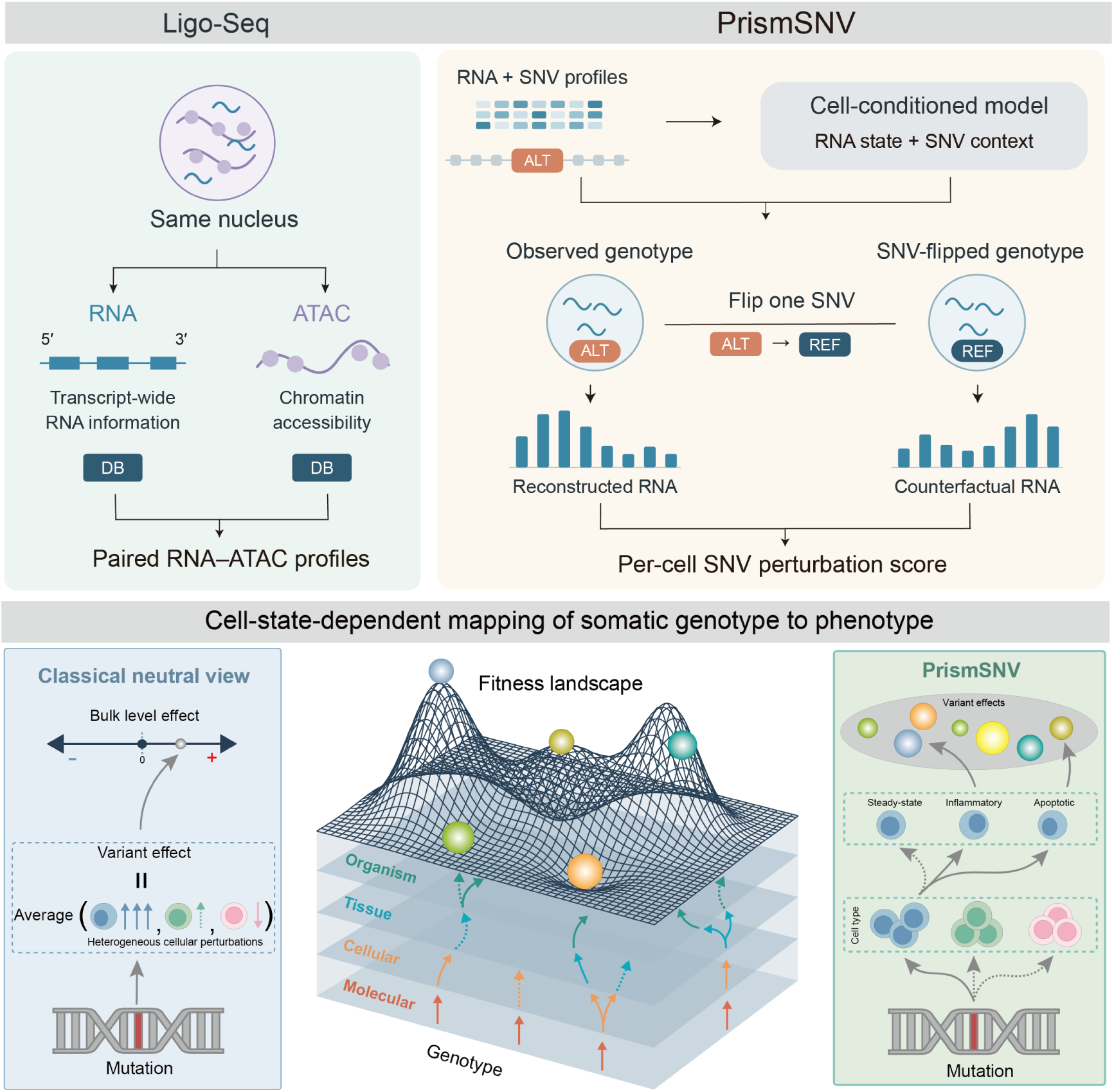

**Highlights:**

- Ligo-Seq jointly profiles transcript-wide RNA and chromatin accessibility
- PrismSNV quantifies transcriptomic effects by toggling SNV states within cells
- Somatic SNV-associated shifts are broadly cell-type-specific across samples
- Less differentiated and transcriptionally plastic states are more sensitive to SNVs

## INTRODUCTION

Cancer genomics has conventionally organized somatic mutations around their consequences for clonal fitness^1–3^. Driver mutations promote tumor progression by conferring selective advantages, whereas passenger mutations are typically defined by their lack of detectable selection at the clone or population level^4,5^. Under this framework, most private^6^, non-recurrent somatic SNVs^7^, which constitute the overwhelming majority of a tumor’s mutational burden^6,8^, are operationally treated as molecularly inert and excluded from mechanistic analyses^1,9^. Whether a variant can be neutral with respect to clonal selection yet produce measurable transcriptomic displacement in specific cellular contexts remains unknown^10^.

This issue reflects a broader gap between increasingly efficient variant detection and functional interpretation^11^. Variants of uncertain significance show that detected variants can influence clinical decisions even when their sequence-annotated pathogenicity remains unresolved, and studies of incomplete penetrance demonstrate that sequence-level pathogenicity does not translate into fixed phenotypic consequence^11–14^. Jensen et al.^15^ showed that genetic modifiers and ascertainment biases contribute to variation in expressivity, highlighting the importance of genetic background in variant interpretation. In cancer genomes, this interpretive gap is amplified by the abundance of private, sparsely observed SNVs^16^ embedded within heterogeneous cellular states^16,17^, such that the molecular effect of any given variant may depend on the specific cell state in which it is expressed.

Existing approaches to variant interpretation capture different, but incomplete, aspects of variant function. Sequence-based tools predict molecular consequence from changes in the reference sequence^18,19^, and population-level analyses infer fitness relevance from recurrence across samples^20–23^; however, these approaches are poorly suited to the private, non-coding or synonymous variants that constitute much of a tumor’s mutational burden^24,25^. Engineered perturbation screens measure transcriptional responses to experimentally introduced variants^26^, but require predefined interventions and matched controls, conditions not met by naturally occurring somatic SNVs that arise endogenously and are sparsely distributed across cells^2,27^. Thus, in most current workflows, variant function is assigned to the mutation itself, as a single context-independent label or score. Yet genetic effects are known to vary across cellular states, as shown by dynamic eQTL effects during CD4^+^ T-cell activation and state-dependent allele-specific expression across T-cell states^28,29^. For endogenous somatic SNVs, the challenge is therefore not simply to annotate the variant, but to quantify its transcriptomic effect within the cellular state in which it naturally occurs.

A further complication is that mutation carriage is not random, cells harboring a given SNV descend from a common ancestral clone and so share genetic background, spatial microenvironment^30^, and often differentiation state^31^. Consequently, unadjusted comparisons between mutation-positive and mutation-negative cells cannot distinguish the contribution of the focal mutation from these correlated cellular and clonal backgrounds^32^. This confounding reflects a structural feature of the variant-generating process in cancer^33^ and is unlikely to be fully resolved by larger sample sizes. Moreover, intertumoral heterogeneity is unlikely to be explained by canonical driver events alone^34,35^, and an expanding body of evidence suggests that passenger mutations are not necessarily molecularly inert. For example, deleterious passenger mutations can impede cancer progression^35^ and the aggregate burden of non-recurrent mutations may shape tumor phenotypes in ways not captured by driver-centric models^36^. Germline-somatic interaction further complicated this landscape, as inherited polymorphisms can modify the functional impact of somatic events in a tissue-specific manner^37^. Bulk-level or prior-driven strategies therefore have limited ability to resolve the perturbation effects of individual somatic variants within their native cellular contexts^38^.

To characterize the molecular effects of private somatic variants across cellular states, we combined Ligo-Seq with PrismSNV (**Figure 1**). Ligo-Seq pairs high-throughput, full-length transcriptome profiling with matched chromatin accessibility, extending RNA coverage across gene bodies to improve the recovery of sparse, endogenous SNVs. PrismSNV then identifies SNVs directly from single-cell reads and learns a batch-adjusted representation of cellular states. For each SNV, PrismSNV compares predicted transcriptomes under alternative allelic states while holding the cellular representation fixed to quantify variant-associated transcriptomic displacement. This framework enables cell-resolved analysis of private, non-recurrent variants without requiring population-level recurrence or experimental perturbation labels.

**Figure 1.**
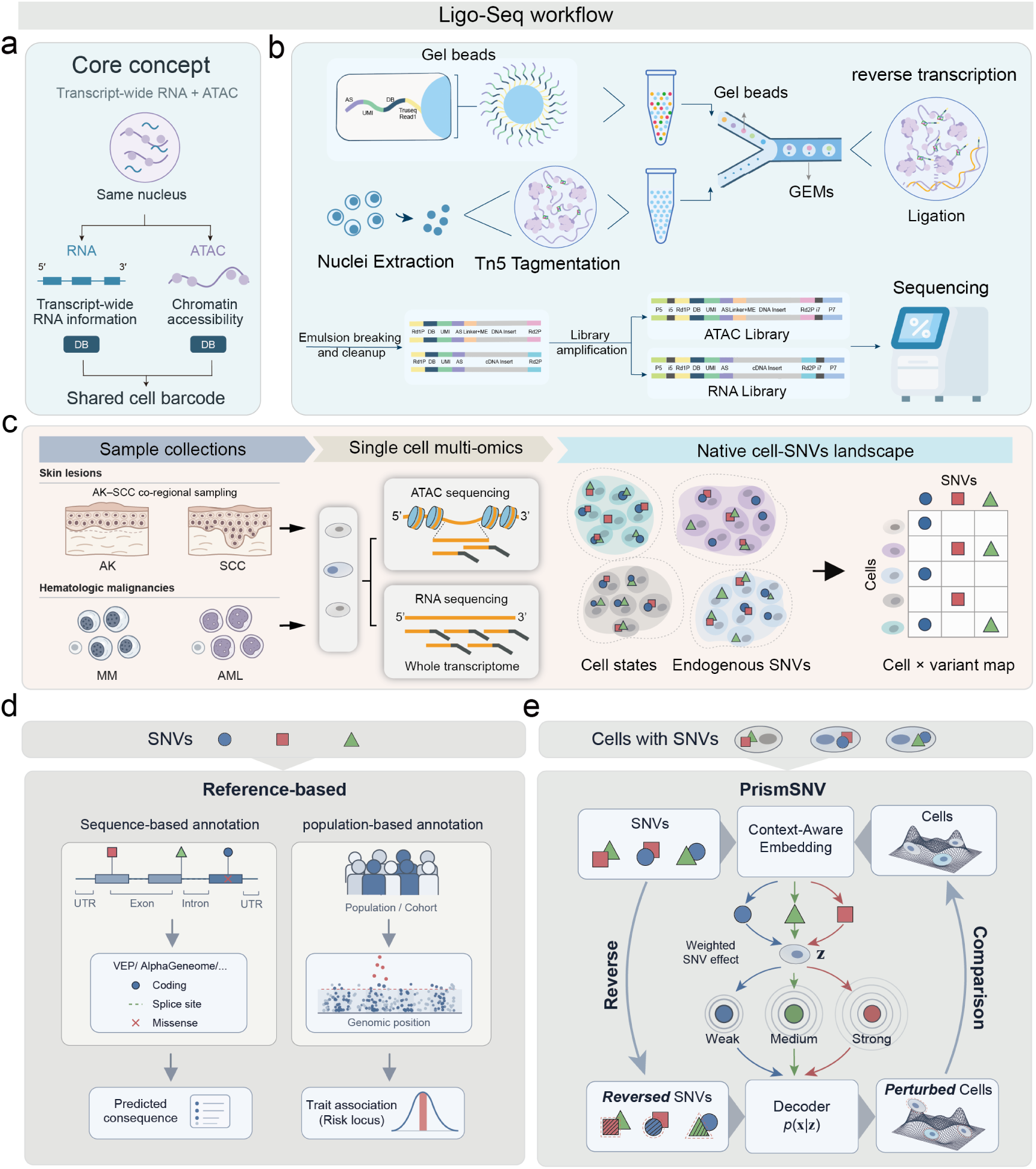
Ligo-Seq and PrismSNV map endogenous somatic SNV-induced transcriptomic displacement effects in native cellular contexts. **a.** Conceptual overview. Ligo-Seq captures RNA sequence information and chromatin accessibility from the same nucleus, linking the two modalities through a shared cell barcode. **b.** Experimental workflow. Following nuclei isolation, accessible chromatin is fragmented and tagged by Tn5 transposase. Nuclei and barcoded gel beads are then co-encapsulated in gel beads-in-emulsion (GEMs) using microfluidics, followed by ligation and reverse transcription. Emulsion breaking, cleanup, and library amplification yield RNA and ATAC libraries for sequencing. Shared cell barcodes enable pairing of the two modalities at single-cell resolution. **c. Left.** Schematic overview of the experimental design and single-cell multi-omics workflow. Ligo-Seq was performed on three different cancer types (including co-regional sampling of AK and cSCC lesions, AML samples and MM samples). Tissues were dissociated, and single cells were subjected to microfluidic partitioning to simultaneously profile chromatin accessibility (ATAC-seq) and whole transcriptomes (full-length) within the same cell**. Right.** Single-cell profiling resolves endogenous somatic SNVs within their native cellular contexts. The resulting cell-by-variant map captures which cells carry which variants, allowing PrismSNV to estimate transcriptomic displacement by comparing observed and counterfactual allelic states within matched cellular backgrounds. **d.** Conceptual distinction between reference-based variant prioritization and context-resolved displacement mapping. Sequence-based tools (VEP, SnpEff, MutationTaster) predict molecular consequence from departure from reference sequence; population-level association infers fitness relevance from recurrence across tumors (driver– passenger classification). **e.** PrismSNV measures transcriptomic displacement—the Euclidean distance between transcriptomes under observed versus counterfactual allelic states—within the native cellular context, independent of external annotation or population data.

We applied this framework to cutaneous squamous cell carcinoma (cSCC), acute myeloid leukemia (AML), and multiple myeloma (MM), observing cell-state-dependent SNV-associated transcriptomic displacement across scRNA-seq platforms and cancer types. Analysis of an independent CD4⁺ T-cell eQTL dataset further revealed variation in the magnitude of genetic regulatory effects during activation. Together, these findings support the variant–cell pair as a unit for interpreting somatic variation, linking variant-associated transcriptomic displacement to cellular identity and state (**Figure S3a**).

## RESULTS

### Ligo-Seq and PrismSNV enable cell-resolved analysis of somatic SNVs

To jointly profile transcriptomes and chromatin accessibility while retaining broader RNA sequence coverage than conventional 3’-capture methods, we developed Ligo-Seq, a high-throughput single-nucleus ATAC-RNA co-assay based on scFAST-seq chemistry. Tagmented nuclei are co-encapsulated with barcoded gel beads, allowing RNA-derived cDNA and accessible chromatin fragments from the same nucleus to acquire a shared cell barcode and unique molecular identifiers. Semi-random 12 + 7 primers replace oligo-dT priming, extending RNA coverage beyond poly(A)-proximal 3’ ends to internal gene-body and untranslated regions that are informative for SNV detection. Following pre-amplification, RNA and ATAC products are separated and sequenced as independently indexed but cell-matched libraries (**Figure 1b**, see Methods for details).

We benchmarked Ligo-Seq against established single-cell RNA and ATAC technologies using mouse brain reference datasets processed with uniform quality-control criteria (**Figures S1a-h**). Across nine scRNA-seq platforms, Ligo-Seq detected a median of 3,994 genes per cell, comparable to 10x Genomics v2, while maintaining low mitochondrial RNA fractions and strong marker-based cell-type separation (**Figures S1a-c**). For chromatin accessibility, Ligo-Seq achieved a median TSS enrichment of 10.4 and a FRiP of 0.65, comparable to TXCI-ATAC and higher than those of the evaluated 10x scATAC and Multiome datasets (**Figures S1e, f**). These benchmarks demonstrate that Ligo-Seq combines sensitive transcript detection with high-quality chromatin-accessibility profiling.

Ligo-Seq provides broader RNA coverage for SNV recovery and matched chromatin-accessibility profiles for regulatory interpretation. In parallel, we developed PrismSNV (**Figures S2 and 1e**), a computational framework for identifying somatic SNVs (**Figures S3b-d**) and quantifying their transcriptomic effects at single-cell resolution. PrismSNV detects candidate SNVs from single-cell transcriptomic reads, applies quality and recurrence-based filtering to distinguish high-confidence somatic variants from sequencing and alignment artefacts, and assigns each retained SNV to the cells in which the alternative allele is observed. PrismSNV learns SNV-informed transcriptomic updates over a fixed RNA-derived cellular representation, using residual regularization and a matched-versus-shuffled reconstruction objective to encourage the use of SNV-specific information. It then quantifies each variant’s associated transcriptomic displacement by toggling its allelic state within the same encoded cellular context (**Figure 1e**, see Methods). PrismSNV accommodates single-cell transcriptomic data with sufficient variant-supporting coverage.

### PrismSNV exposes widespread somatic perturbations at single-cell resolution

To characterize somatic SNV-associated transcriptomic displacement across distinct cancer contexts, we applied PrismSNV to 26 samples spanning cSCC, acute myeloid leukemia (AML), and multiple myeloma (MM) (**Figure 1c**). The skin cohort comprised nine matched actinic keratosis (AK)-cSCC pairs and one additional cSCC lesion from ten patients, encompassing diverse cellular states across premalignant and malignant lesions. Histopathological examination confirmed lesion identity, and single-cell RNA and ATAC analyses resolved the major skin cell populations (**Figures S4d-g**). The hematological cohorts comprised bone marrow samples from three patients with AML and four patients with MM. RNA marker genes, ATAC gene activity markers, and associated cell-type annotations for all three cohorts are provided in **Supplementary File 1**. Transcriptomic and chromatin accessibility measurements were obtained from the same individual cells (**Figure 1a, b**).

PrismSNV integrates batch correction and representation learning with probabilistic SNV genotyping to characterize variant-associated transcriptomic displacement in individual cells. We use the AK-cSCC cohort as an example. For this cohort, batch correction and representation learning incorporated public human skin datasets^39–41^ comprising 711,213 cells, together with the newly generated AK-cSCC datasets (**Figure S5a**). Probabilistic allelic states were assigned to candidate somatic SNVs while accounting for sequencing errors, allelic dropout, and sample-specific allele frequencies (**Figures S3b-d**), with the distribution of detected SNVs summarized in Figure S4a. We systematically evaluated pretraining performance (**Figure S5a**), computational metrics (**Figures S5b and S5c**), the filtering strategy (**Figure S3c**), sampling-depth-associated errors (**Figures S5f and S5g**), permutation-based assessment of perturbation scores (**Figure S5h**), the false-positive rate (**Figure S5i**), and potential bias associated with cell-type-specific marker genes (**Figures S5j and S5k**). Attention scores were positively correlated with final SNV perturbation scores (**Figures S5d**, **e**). Following these evaluations, we ranked cell type-SNV pairs by attention score and retained the top 10,000 pairs. Deduplication across cell types yielded 4,617 unique candidate SNVs for downstream analysis in the AK-cSCC cohort.

Comparing SNVs prioritized by PrismSNV against those selected by sequence-based annotations revealed a striking divergence in their transcriptomic impact (**Figure S1i-l**). Each method selected candidates according to its own predefined criterion: PrismSNV candidates comprised the top 10% of SNVs ranked by decoder-based counterfactual Euclidean displacement within each cell type, whereas VEP, SnpEff and MutationTaster candidates were selected using their respective HIGH-impact or Deleterious classifications (**Figure S1i**). Candidate selection was completed without reference to the subsequent RNA distance-delta metric. We then evaluated all method-prioritized candidate sets using the same RNA-based endpoint. For each SNV, RNA distance delta was defined as the observed Euclidean distance between the mean transcriptomes of ALT- and REF-supported cells minus the mean distance obtained after randomized cell-group assignments with preserved group sizes. Random background distributions were generated by repeatedly sampling eligible SNVs from the same candidate universe. Variants prioritized as deleterious or high impact by sequence-based annotation methods showed RNA distance deltas comparable to the random-SNV background (two-sided Wilcoxon rank-sum test, *P* = 0.36 ; Cohen’s *d* = −0.22 ; **Figure S1i, j** and **Supplementary File 2**). In contrast, PrismSNV-prioritized SNVs showed substantially greater RNA distance deltas than matched random-SNV sets (two-sided Wilcoxon signed-rank test, *P* = 0.0049; paired Cohen’s *d_z_* = 0.93; **Figure S1l** and **Supplementary File 2**).

PrismSNV further revealed pronounced heterogeneity in perturbation magnitude across AK-cSCC cell populations, with mean perturbation-distance deltas varying 23.5-fold across cell types (range, 9.8-230.6; median, 78.1; IQR, 56.6-108.6), and background-standardized enrichment spanning Z = 0.43-9.92 (median Z = 6.03; IQR, 3.71-7.66; **Supplementary File 2**). Similar heterogeneity was observed in AML and MM, with mean perturbation-distance deltas varying 10.6-fold across five evaluable AML cell populations (range, 3.4-36.4; median, 18.7; IQR, 14.1-21.3) and 25.1-fold across nine evaluable MM cell populations (range, 5.7-143.2; median, 21.4; IQR, 11.9-49.3; **Supplementary File 2**). Individual variants produced strong transcriptomic displacements in some cellular states while remaining near background levels in others (**Figure S5h and S7d**). This pattern indicates that the transcriptomic displacement produced by a somatic variant is not uniform across cellular contexts, but varies with the molecular state of the cell^33^.

### Cell-type specificity is the dominant mode of somatic variant action

To quantify the extent of cell-type dependency, we evaluated all eligible SNVs across diverse cell populations (**Figure 2a**). We evaluated 4,617 candidate SNVs selected based on the model’s top-ranked SNV attention scores. To assess cell-type dependence, we quantified η² for top 30,000 prespecified high-coverage candidate SNVs in the AK– cSCC cohort and evaluated significance using 1,000 permutations of cell-type labels within host samples, followed by Benjamini–Hochberg correction. The observed η² distribution was markedly shifted towards larger values relative to the SNV-specific shuffled-label null (**Figure 2b**). Among SNVs with empirical BH-FDR *q* < 0.05, 25,021 (83% of SNVs) exceeded the 10% relative-amplitude threshold and 19,828 (66%) exceeded the 30% threshold, a similar pattern was also evident in AML and MM samples (**Figure S6a,b**), extending this observation to hematological malignancies, demonstrating that pronounced cell-type-specific transcriptomic displacement is widespread among candidate SNVs.

**Figure 2.**
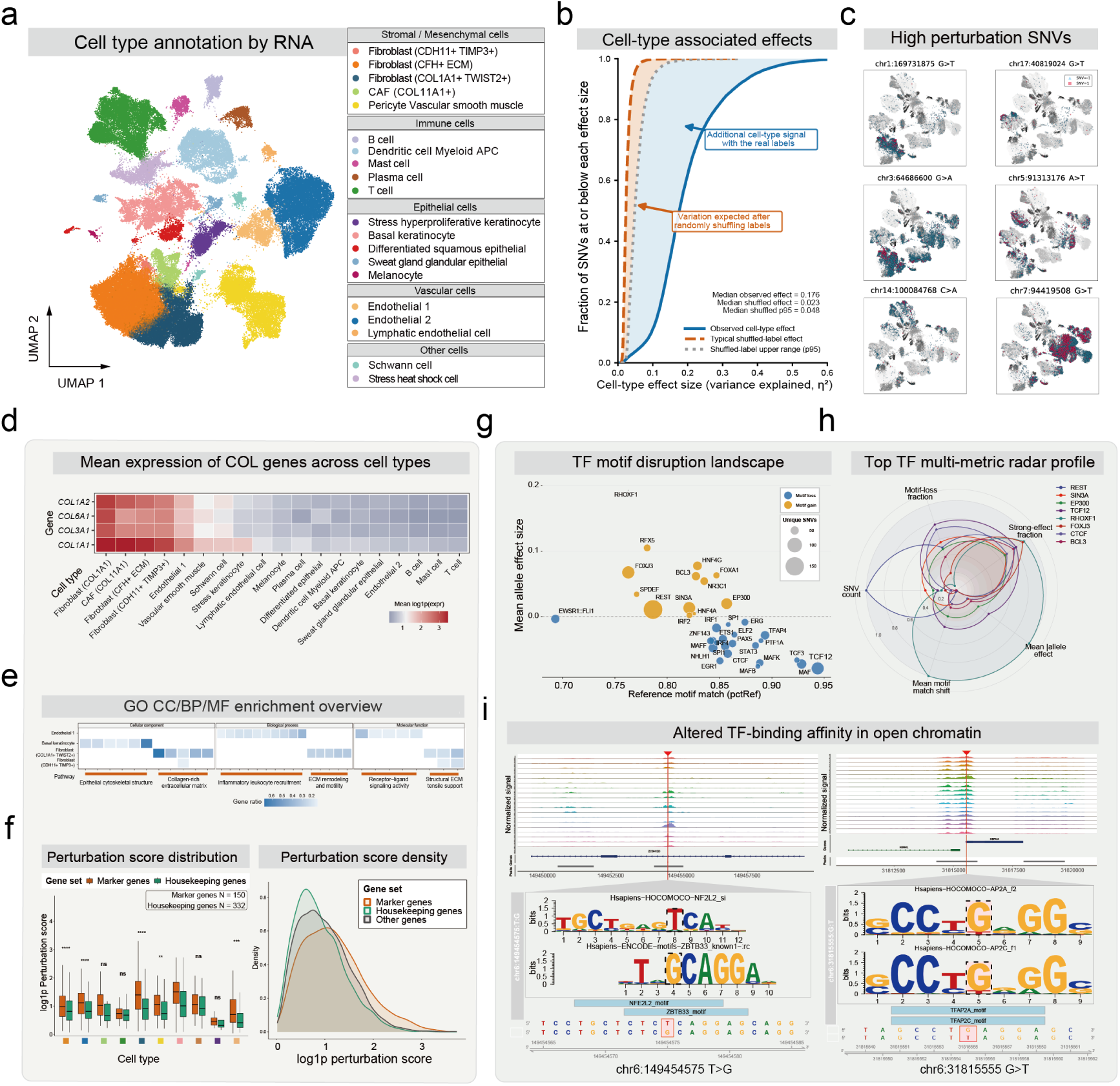
Cell-type-specific transcriptomic displacement of somatic SNVs. **a.** UMAP projection of full-length scRNA-seq data from all full-length samples (9 samples, including 5 cSCC and 4 AK), colored by annotated lineage. **b.** Empirical cumulative distribution functions show cell-type effect sizes (η²) across 30,000 coverage-defined candidate SNVs. Cell-type labels were permuted 1,000 times within host samples. The observed η² distribution (blue) was shifted towards larger values than the SNV-specific permutation median (orange dashed) and null 95th percentile (grey dotted). The median observed η² was 0.176, compared with a median shuffled-label value of 0.023. Overall, 28,644 SNVs (95 %) exceeded their own null 95th percentile, and 28,579 (95 %) were significant after empirical BH-FDR correction (*q* < 0.05), demonstrating substantially stronger cell-type-associated displacement than expected from random label shuffling. **c.** UMAP visualizations of single-cell transcriptomic profiles for representative high-perturbation SNVs. Cells are colored by genotype: mutant allele present (red, SNV=1), unmutated/reference (blue, SNV=-1), and undetected/uncertain (gray). Spatial segregation between the mutant and reference populations reflects variant-associated transcriptomic displacement. **d.** Expression patterns of major collagen family genes across different cell types. **e.** Gene Ontology (GO) enrichment overview (CC: **C**ellular **C**omponent, BP: **B**iological **P**rocess, MF: **M**olecular **F**unction) for genes harboring high-perturbation SNVs. Significant enrichments (calculated by ClusterProfiler with default parameter, adjust *P* < 0.05) observed in endothelial cells (angiogenesis), basal keratinocytes (epidermal differentiation), and fibroblasts (extracellular matrix organization). **f. Left**, SNVs occurring in housekeeping genes exhibited significantly (Mann-Whitney U test, ns, p >= 0.05; *, p < 0.05; **, p < 0.01; ***, p < 0.001; ****, p < 0.0001.) lower perturbative effects compared to those in cell-type-specific genes, as well as compared to background SNVs. **Right**, density plot compares the distribution of log1p-transformed SNV perturbation scores across marker genes, housekeeping genes, and other genes; marker-gene SNVs show a right-shifted distribution, suggesting stronger perturbation effects overall. **g.** TF motif effect landscape summarizing motifbreakR predictions. The x axis shows reference-allele motif matching scores (pctRef), and the y axis shows mean allelic effect sizes for the indicated TF motifs. Point size represents the number of associated unique SNVs. Orange and blue indicate positive and negative mean allelic effects, corresponding to predicted motif gain and loss, respectively. **h.** Radar profiles of eight selected TFs showing motif-loss fraction, strong-effect fraction, mean allelic effect size, mean motif-match shift, and the number of associated SNVs. Each colored profile represents one TF. **i.** The multi-omic validation of regulatory SNVs. Paired scATAC-seq tracks (**top**) confirm these loci reside in open chromatin regions across various cell lineages (calculated by SnapATAC2), different tracks mean different cell types, peaks mean open chromatin regions. Motif analysis (**bottom**) reveals that these specific single-nucleotide alterations (highlighted in red box) critically disrupt transcription factor binding affinities (calculated by motifbreakR).

Conventional variant annotation tools did not identify the same cell-resolved high-displacement variants (**Figure S4b, c and S7e**). VEP impact rank was essentially uncorrelated with PrismSNV perturbation score (Spearman ρ = −0.038). Among 606 evaluable SNVs annotated as HIGH impact by VEP, 40.8% (247/606) showed no detectable single-cell perturbation in PrismSNV, with perturbation scores falling within the matched background distribution. Conversely, among PrismSNV-identified SNVs with transcriptomic displacement significantly above background, 69.6% (284/408) were not captured by any of the three conventional annotation tools (one-sided exact binomial test against an equal captured/uncaptured expectation, *P* = 7.6 × 10⁻¹⁶). Thus, sequence-based pathogenicity annotations were poorly aligned with cell-resolved transcriptomic displacement (**Figure 2c**). These results support the variant-cell pair, rather than the variant alone, as the relevant unit for measuring somatic variant impact.

### Private somatic variants produce cell-state-specific transcriptional displacement

Given that variant effects are primarily cell-type-specific, we next examined whether private somatic variants, which constitute the majority of the mutational burden and are largely absent from clinical databases, produce detectable molecular phenotypes. Overall, 54 of 2,123 high-perturbation SNVs pooled across the three cohorts were catalogued in ClinVar^42^ (2.54%; AK–cSCC, 22/713; AML, 9/596; MM, 23/814), and AlphaGenome produced broad perturbation predictions (median, 360 scRNA-seq predictions per SNV; mean, 439.4) even when evaluated in skin-relevant tissue contexts (**Figure S7c**). Across the retained skin-related RNA predictions, individual SNVs showed broad distributions of absolute quantile scores (**Figures S7a, b**). In contrast, the cells harboring high-perturbation SNVs identified by PrismSNV showed transcriptomic displacement relative to their non-mutant counterparts (**Figure S1l; Figure S7c-d**), indicating that private somatic variants generate measurable molecular phenotypes when evaluated in their native cellular context. These displacement patterns were associated with two non-mutually exclusive features. One pattern involved SNVs located in genes embedded in cell-type-specialized programs. Although genes harboring all candidate SNVs did not show significant pathway enrichment after multiple-testing correction, genes carrying high-perturbation SNVs, defined as the top 10% of candidate SNVs by perturbation score within each cell type, showed significant enrichment for cytoskeletal organization, inflammatory response, and cell migration programs in fibroblast subtypes, basal keratinocyte cells and endothelial cells (clusterProfiler, adjusted *P* < 0.05; **Figure 2e**). For example, somatic SNVs in collagen family genes (*COL12A1*: chr6:75191884:G>T and *COL1A1*: chr17:50188982:C>G) exerted substantially stronger perturbative effects in fibroblasts than in other cell types, consistent with the central role of collagen genes in extracellular matrix organization in fibroblasts (**Figure 2d**). Similar context-matched perturbation patterns were also observed in other cell types (**Supplementary File 3**). Notably, high expression alone did not predict SNV sensitivity. Housekeeping genes^43^ (n = 2,158), despite being broadly and highly expressed across nearly all cell types, showed lower SNV-associated perturbative effects than cell-type-specific genes or background SNVs (**Figure 2f**). This indicates that cellular-context dependency, rather than expression level, is associated with variant transcriptomic impact.

A second pattern involved predicted disruption of accessible cis-regulatory elements. We retained the top 30% of SNVs within each cell type ranked by cosine-distance perturbation score, yielding 1,862 unique SNVs after deduplication across cell types. MotifbreakR returned predictions for 1,748 of these SNVs. Integration with matched ATAC-seq profiles identified 332 selected SNVs overlapping accessible chromatin regions, of which 302 exhibited strong predicted effects on transcription-factor motifs. Among these 302 SNVs, 87 showed motif gains only, 74 showed motif losses only, and 141 showed gains for some motifs and losses for others (**Supplementary File 4**).

These 302 candidate regulatory SNVs were distributed across 264 ATAC-seq peaks (**Supplementary File 4**). To link these candidate regulatory elements to putative target genes, we considered all peak-gene pairs in which the transcription start site lay within 500 kb upstream or downstream of the candidate peak. For each pair, we calculated the Pearson correlation coefficient (r) between peak accessibility and gene expression across cells. Correlation z-scores and nominal P values were estimated using background peaks matched for GC content, mean accessibility, and peak length. Under an initial threshold of |*r*| ≥ 0.05 and nominal P < 0.05, 132 ATAC-seq peaks harboring 158 SNVs (52.3%) were associated with the expression of at least one gene, yielding 273 distinct peak-gene links involving 249 genes. Applying more stringent criteria of |*r*| ≥ 0.1and nominal P < 0.01 retained 78 SNVs within 60 ATAC-seq peaks, corresponding to 81 peak-gene links and 78 candidate target genes.

Notably, a single SNV could affect motifs for multiple transcription factors simultaneously (**Figure 2h, i, Figure S8a, d**). These SNV-affected motifs were enriched in regulatory programs related to stemness, lineage specification and immune/inflammatory transcriptional control (**Figure S8b, c**). Because transcription factors are differentially expressed across cell types and states within the same tissue (**Figure S8e**), the regulatory consequence of a given SNV is likely to depend on the cellular context in which it occurs, providing a plausible mechanism for cell-to-cell heterogeneity in SNV-induced perturbation.

Together, these analyses show that private somatic SNVs can produce measurable transcriptomic displacement in specific cellular contexts, even when they lack clinical annotation or recurrence-based support. Variants that would be deprioritized under conventional driver-passenger frameworks may nevertheless perturb gene-expression states in the cells in which they occur.

### Experimental validation confirms the impact of high-perturbation SNVs

Most high-displacement somatic SNVs prioritized by PrismSNV were restricted to individual patients in our cohort, limiting their evaluation through conventional variant-level association analyses. To assess their functional relevance experimentally, we focused on candidates prioritized in the fibroblast compartment of cutaneous squamous cell carcinoma (cSCC) and selected two high-displacement variants in the extracellular matrix gene *COL3A1*, p.G726E and p.G858D, for in vitro characterization.

We used a knockdown-rescue strategy in human skin fibroblasts (HSFs) to evaluate the effects of the COL3A1 variants (**Figure 3a**). Following COL3A1 knockdown, cells were rescued with WT COL3A1, G726E or G858D, with an empty-vector control included for comparison. Construct sequences and protein expression were assessed by Sanger sequencing and western blotting (**Figures 3b-d**).

**Figure 3.**
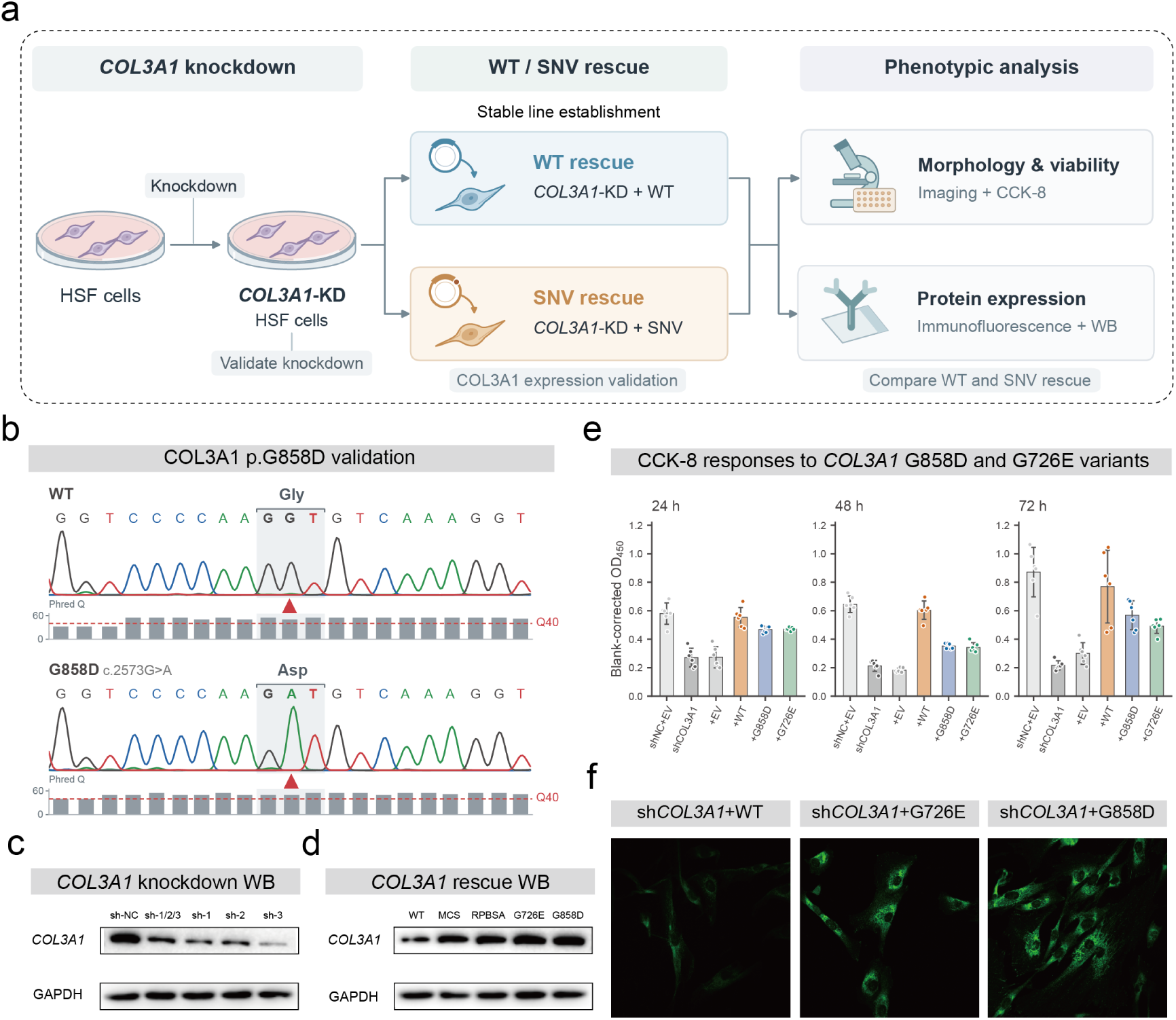
Experimental validation of COL3A1 variants with strong predicted perturbation effects. **a.** Schematic of the experimental workflow. Following *COL3A1* knockdown in human skin fibroblasts (HSFs), stable cell lines expressing wild-type (WT) *COL3A1* or the G726E or G858D variants were established. Cellular phenotypes and protein expression were assessed by microscopy, CCK-8 assays, immunofluorescence, and western blotting. **b.** Sanger sequencing chromatograms confirming the COL3A1 rescue constructs. Sequencing verified the reference coding sequence in the WT vector and the intended G858D substitutions in the mutant vectors. **c.** Western blot analysis of HSFs expressing control shRNA (sh-NC), sh-1, sh-2, sh-3, or sh-1/2/3 targeting *COL3A1*. The upper and lower panels show *COL3A1* and the loading control, respectively. **d.** Western blot analysis of *COL3A1*-knockdown cells (**c**) rescued with wild-type *COL3A1* or the G726E and G858D mutants. The upper and lower panels show *COL3A1* and the loading control, respectively. **e.** CCK-8 measurements at 24, 48, and 72 h in control cells (shNC+EV), *COL3A1* knockdown cells (sh*COL3A1*), and knockdown cells expressing empty vector (EV), WT *COL3A1*, G858D, or G726E. Results are presented as blank-corrected absorbance at 450 nm (OD₄₅₀). Bars represent the mean ± SD of six technical replicate wells per group at each time point. Each dot represents one technical replicate well. **f.** Representative confocal fluorescence images of *COL3A1* knockdown HSFs expressing WT *COL3A1*, G726E, or G858D, showing the distribution of the green fluorescence signal.

Upon *COL3A1* knockdown, the wild-type (WT) rescue group exhibited higher mean CCK-8 signal compared with the empty vector (EV) control group. Under uniform seeding density and culture conditions (six replicate wells per group), background-subtracted OD_450_ values at 24, 48, and 72 h were consistently lower in both G858D and G726E mutant rescue groups than in the WT rescue group (**Figure 3e**). This reduction in CCK-8 responsiveness indicates that these mutations may attenuate cellular proliferation and metabolic reductive capacity relative to WT.

HSF cells expressing WT *COL3A1* or the G726E or G858D variant following *COL3A1* knockdown were cultured under identical conditions, immunostained for COL3A1, and imaged by confocal microscopy using identical acquisition settings. Representative images showed stronger intracellular *COL3A1* fluorescence in the G726E and G858D rescue groups than in the WT rescue group (**Figure 3f**), suggesting that these mutations may impair normal *COL3A1* secretion and lead to intracellular retention.

### Differentiation state modulates transcriptomic displacement magnitude

We next investigated whether sensitivity to SNV-induced perturbation varies across cellular states within the same lineages. By aggregating SNV-level perturbation scores at the single-cell level and normalizing by the expression level to exclude sequencing depth confounding (**Figure S5f, g**), we projected the resulting sensitivity landscape onto the global UMAP space (**Figure 4a**). This revealed substantial heterogeneity not only across lineages but also within individual lineages. Major skin-resident populations, including keratinocytes, fibroblasts and endothelial cells, displayed broad intra-cluster variation in perturbation sensitivity (**Figure 4b**).

**Figure 4.**
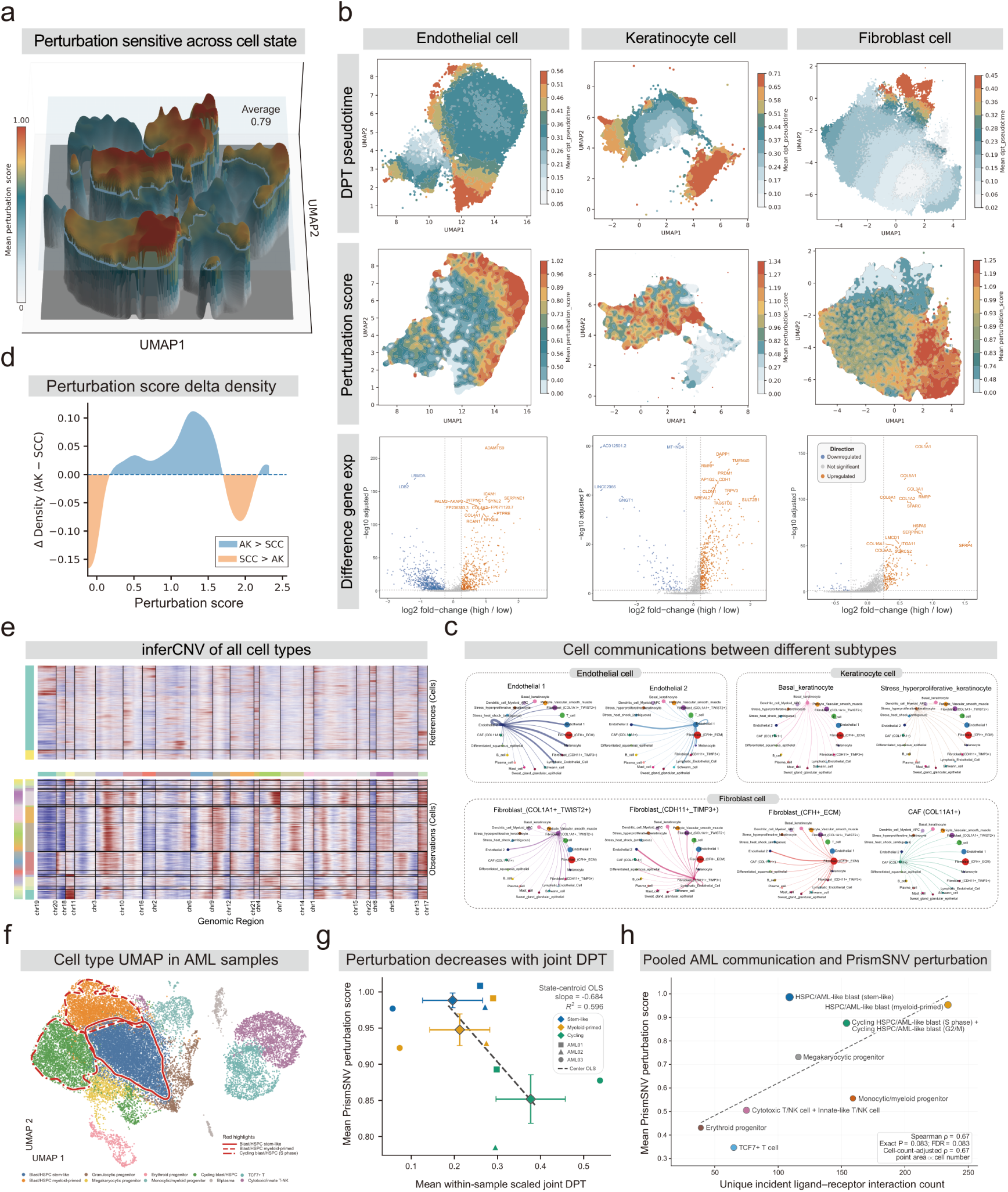
Cell state modulates transcriptomic displacement magnitude. **a.** 3D landscape of displacement magnitude across cSCC cells after Gaussian smoothing. z axis: normalized displacement score (0-1); blue plane: mean displacement level (0.79). **b.** Top row: Pseudotime trajectories for endothelial cells, keratinocytes and fibroblasts. Middle row: Displacement magnitude mapped onto pseudotime, showing inverse association with differentiation. Bottom row: Differential expression between top and bottom quartiles of displacement magnitude: highly sensitive cells upregulate inflammatory activation, ECM remodeling and tumor stromal programs. **c.** CellChat-inferred cell-cell communication networks for endothelial, keratinocyte and fibroblast subtypes. Each network shows signaling connections associated with the indicated subtype. Nodes represent cell populations, and edges represent inferred intercellular communication links. **d.** Paired AK–cSCC comparison (n = 4 patients, full-length sequencing). Delta density plot: AK cells show globally higher displacement magnitude than matched cSCC cells. Blue: Density AK > Density cSCC; orange: Density cSCC > Density AK. **e.** inferCNV analysis across all cell types. Heatmaps display large-scale chromosomal copy number variation patterns for reference (top, T cell and B cell) and observation cells (bottom, other cell types). Right histogram shows standardized expression scores. Clear differences in copy number alteration burdens are visible between distinct cell subsets. **f.** UMAP visualization of single-cell profiles from AML samples. The red circle delineates the hematopoietic stem and progenitor cell (HSPC) compartment. **g.** Scatter plot depicting the relationship between cellular perturbation scores and diffusion pseudotime (DPT) within HSPCs. Across three independent patient samples, HSPC perturbation scores consistently decline along the DPT trajectory. **h.** Correlation analysis between cellular perturbation scores and intercellular communication strength (quantified by the number of inferred ligand-receptor interactions) across all cell types, including HSPC subpopulations, in AML samples. A positive correlation is observed between these two metrics.

To determine whether intra-lineage variation in SNV perturbation sensitivity reflected distinct cell states, we examined fibroblasts, keratinocytes and endothelial cells along their inferred trajectories (**Figure 4b**). In AK-cSCC samples, perturbation sensitivity was inversely associated with pseudotime within each lineage; SNV-induced perturbation was strongest in transcriptionally plastic states and declined progressively toward committed, malignant fates. Differential expression analysis between the top and bottom quartiles of PrismSNV-defined sensitivity revealed that highly sensitive cells preferentially upregulated programs related to inflammatory activation, extracellular matrix remodeling and tumor stromal activation (**Figure 4b, bottom**). Cell-cell communication analysis further showed stronger signaling activity in highly sensitive cells (**Figure 4c**), indicating that SNV effects are amplified in transcriptionally plastic and functionally interactive cell states. Across 4 patients with paired AK-cSCC lesions, cells from premalignant AK showed systematically higher perturbation scores than cells from matched malignant cSCC lesions (**Figure 4d**). inferCNV analysis revealed distinct large-scale copy-number patterns across these cellular subsets (**Figure 4e**).

Furthermore, intra-lineage variation in perturbation magnitude was also detected in hematological malignancies, including three AML patient samples (Figures 4f and 4g) and four MM patient samples (**Figures S6c-e**). Specifically, within the AML hematopoietic stem and progenitor cell (HSPC) compartment (**Figure 4f**, red circle), the mean cellular perturbation score correlated inversely with pseudotime (**Figure 4g**). In parallel, cells actively engaged in robust cell-cell communication were more vulnerable to SNV-induced perturbations (**Figure 4h**). These results support that the transcriptomic displacement by somatic variation is not fixed within a cell type, but varies according to the intrinsic differentiation and malignant state of individual cells.

### State-dependent architecture is conserved across platforms and cancer types

To test whether gene-level SNV displacement profiles are reproducible across patients, distinct private variants, and sequencing platforms, we deployed conventional 3’ scRNA-seq (5 patients, 10 samples) as an orthogonal technical control against full-length scRNA-seq data (5 patients, 9 samples) across shared cSCC cell populations.

Relative to scRNA-seq, Ligo-Seq achieved a roughly fourfold higher fraction of candidate SNVs within coding sequence regions (CDS) (**Figure 5a**). In addition, sequencing-depth-normalized candidate SNV yields were 5.6-fold higher in AK and 9.7-fold higher in cSCC with full-length sequencing than with 3’ scRNA-seq (**Figure 5b**), alongside more balanced coverage across the entire gene body (**Figure 5c**). Despite these platform-level performance differences and Ligo-Seq’s superior capture efficiency, the underlying biological signals remained highly consistent across platforms. At the single-cell level, perturbation scores showed highly concordant cell-type-associated distributions between technologies after excluding cells with no detectable perturbation signal. Among the major shared populations, keratinocytes, fibroblasts, myeloid/dendritic cells, and perivascular smooth muscle cells carried the strongest perturbation burdens in both datasets (**Figure 5d**). We assessed reproducibility by comparing cell-type-level perturbation rankings rather than paired cells. This revealed a substantially conserved perturbation hierarchy across shared cell classes, accompanied by cross-platform concordance in cell-type-level perturbation scores (**Figure 5d middle**). Concordance was further enhanced after aggregating perturbation effects at the gene level, where mean-effect ranks were highly correlated between 3’ and full-length scRNA-seq profiles (**Figure 5d right**; **Supplementary Files 3**, **6**).

**Figure 5.**
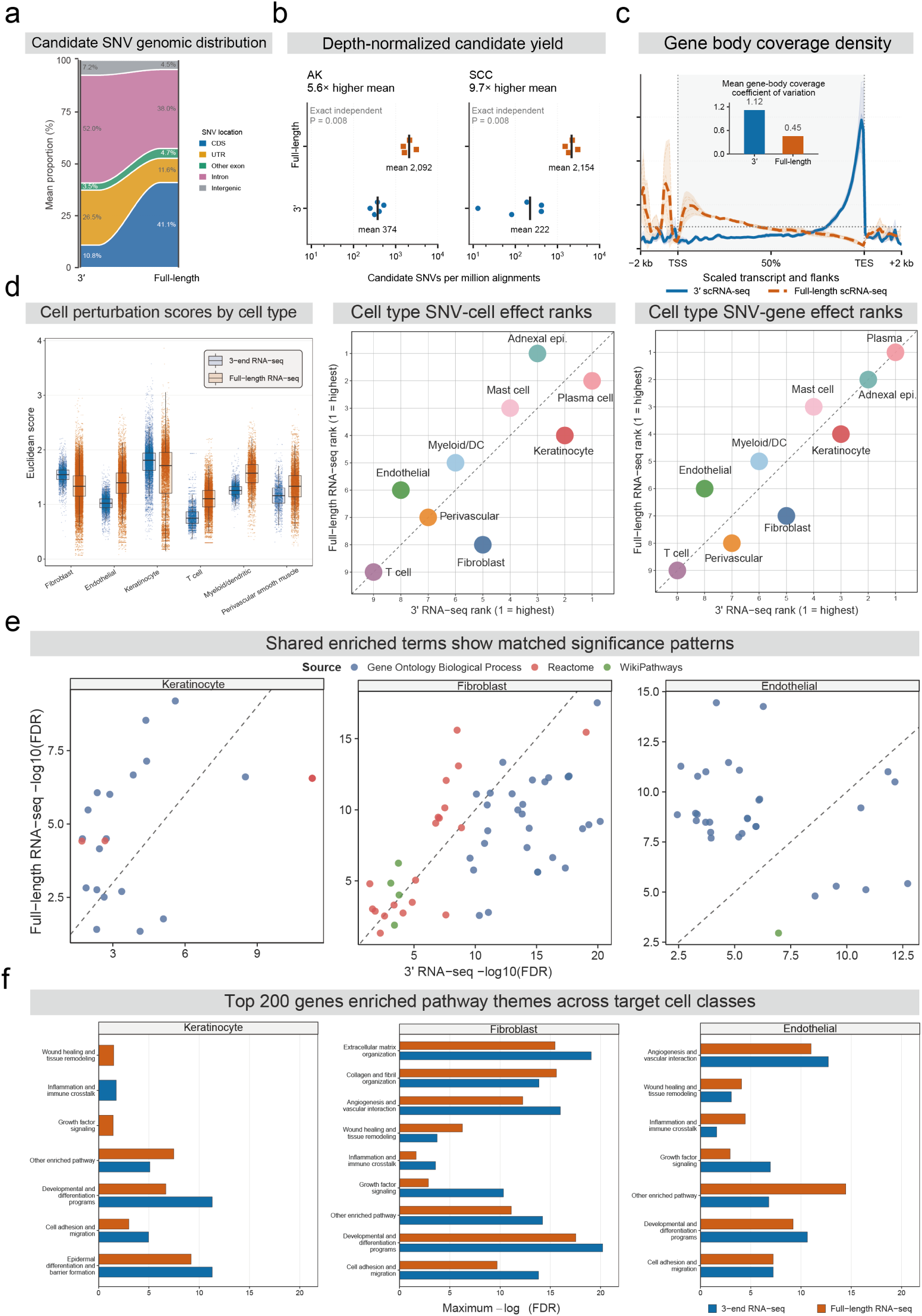
Concordance of gene-level displacement profiles across patients, distinct private variants, and platforms. **a.** Genomic distribution of candidate SNVs from 3’-end and full-length scRNA-seq. Filtered SNVs (AF <20%, excluding RNA-editing sites) were hierarchically categorized (GRCh38: CDS, UTR, other exon, intron, intergenic). **b.** Sequencing depth-normalized candidate SNV yields. In AK and SCC, full-length sequencing (n = 4) showed a 5.6-fold and 9.7-fold increase in mean candidate SNV yield compared with 3’-end sequencing (n = 5), respectively (exact independent permutation test, both P = 0.008). Points represent independent samples; black vertical bars indicate mean values. **c.** Normalized coverage across transcripts and flanking regions. Thick lines indicate sample means, shaded bands represent 95% confidence intervals, and thin lines denote individual samples. Compared with 3’ scRNA-seq (n = 5), full-length scRNA-seq (n = 4) exhibited lower coefficients of variation for gene body coverage across both AK and SCC, indicating a more uniform coverage distribution. **d. Left.** Distribution of transcriptomic displacement score calculated by PrismSNV across shared cell types detected in 3’ RNA-seq (5 patients, 10 samples) and full-length RNA-seq (5 patients, 9 samples). Boxplots: center line = median, box = IQR, whiskers = 1.5× IQR; points = individual cells. **Middle**. Rank concordance of cell-type perturbation scores between full-length and 3’ RNA-seq platforms with 14313 genes, based on median cell perturbation scores aggregated by nine cell types. Each point represents one shared cell type. **Right**. Rank concordance of SNV-gene perturbation effects between full-length and 3’ RNA-seq platforms with 14313 genes, based on median SNV/gene effect scores aggregated by nine cell types. Each point represents one shared cell type. **e.** Cross-platform comparison of shared enriched pathways among top-ranked perturbation-associated genes from 14313 genes in keratinocytes, fibroblasts and endothelial cells. Axes show enrichment significance in each platform, measured as −log10(FDR). Points are shared enriched terms colored by pathway source. **f.** Summary of enriched pathway themes among the top 200 displacement-associated genes from 14313 genes for keratinocytes, fibroblasts and endothelial cells in each platform. Bar length indicates the strongest enrichment significance within each pathway.

We next asked whether this cross-platform concordance also extended to biological interpretation. Enrichment analysis of top perturbation-associated genes (top 10% per cell type) revealed quantifiably matched functional signatures between 3’ and full-length scRNA-seq in keratinocytes, fibroblasts and endothelial cells, despite differences in transcript coverage and variant detectability (**Figure 5e**). Pathway-theme aggregation further resolved coherent cell-type-specific programs in both datasets, including epidermal differentiation and barrier formation in keratinocytes, extracellular matrix organization in fibroblasts, and angiogenic, vascular-interaction and adhesion-related programs in endothelial cells (**Figure 5f**). Thus, even with limited overlap in exact SNVs, PrismSNV retained the dominant biological structure of perturbation effects across distinct scRNA-seq platforms.

PrismSNV was further applied to publicly available 3’ scRNA-seq datasets from intrahepatic cholangiocarcinoma (8 samples, 62,326 cells)^44^, bladder cancer (8 samples, 33,541 cells)^45^ and breast cancer (20 samples, 120,519 cells)^34^ (**Supplementary File 5**). Across distinct tumor tissues, individual SNVs exerted cell-type-specific effects, and different cell types also showed unequal sensitivity to SNV-induced perturbations (**Figure S9a, b**).

The partially conserved perturbation hierarchy and matched functional signatures across distinct platforms and cancer types support that state-dependent genotype-phenotype mapping is not merely a technical artifact specific to cSCC or full-length scRNA-seq.

### Cell-state transitions reshape variant effects during T-cell activation

To examine how genetic effects change during normal cellular activation, we reanalyzed published cis-eQTL summary statistics from a study profiling 655,349 CD4⁺ T cells during anti-CD3/CD28 stimulation^46^. Across naive (TN), central-memory (TCM), effector-memory (TEM), and CD45RA-re-expressing effector-memory (TEMRA) populations, we identified 14 genes with effect-coefficient differences across activation stages and five with differences between populations at matched times. These sets comprised 17 distinct genes, whose representative SNV-gene associations displayed diverse patterns of effect magnitude and direction across cellular contexts (**Figure 6a**).

**Figure 6.**
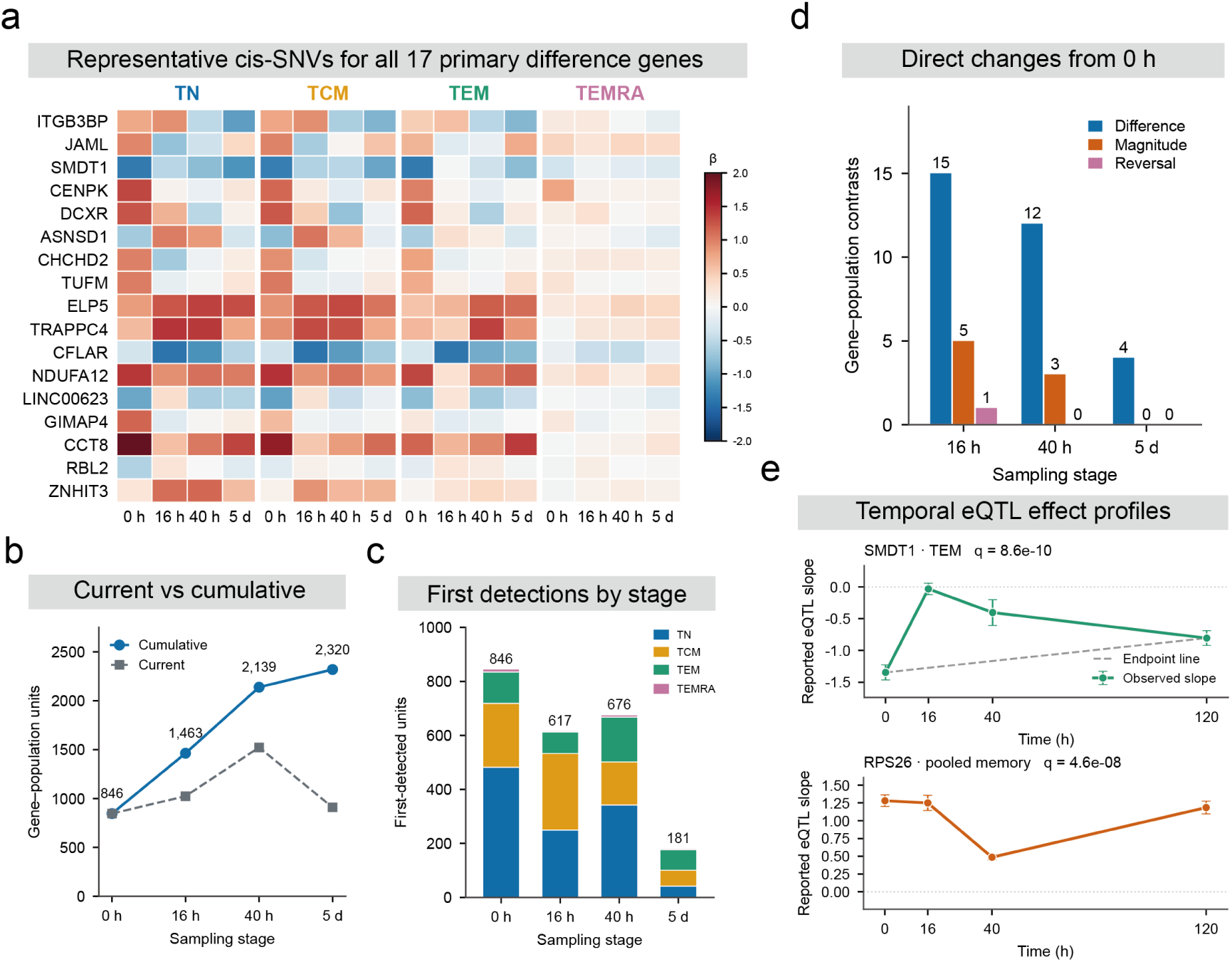
Context-dependent eQTL effects during CD4⁺ T cell activation. **a.** Reported eQTL coefficients (β) for representative cis-SNVs associated with 17 genes showing significant coefficient differences across time within populations or between populations at matched times. Each row represents one fixed SNV–gene pair across naive (TN), central-memory (TCM), effector-memory (TEM), and CD45RA-re-expressing effector-memory (TEMRA) T cells. Colors indicate signed coefficients. Genes were identified by gene-wise Bonferroni correction across eligible SNVs and comparisons, followed by Benjamini–Yekutieli (BY) correction at q ≤ 0.025 in each primary comparison family. **b.** Current and cumulative association detections within a fixed universe of 37,173 gene–population units evaluated across all four sampling stages. Current counts denote units detected at each stage; cumulative counts denote units detected at least once up to that stage. Associations were identified by within-gene cis-SNV Bonferroni correction followed by BY correction across gene–population–time tests (q ≤ 0.05). Cumulative counts are descriptive unions of these discoveries and do not establish the biological onset of an effect. **c.** Numbers of gene–population units first meeting the association criterion in (B) at each sampling stage, stratified by T cell population. Previously detected units were not counted again upon re-detection. **d.** Numbers of gene–population contrasts showing significant differences from the unstimulated condition in signed coefficients (Difference), absolute coefficient magnitudes (Magnitude), or association directions (Reversal). Each later stage was compared directly with 0 h. Conservative summary-statistic tests were followed by within-gene cis-SNV Bonferroni correction and BY correction across gene– population–contrast tests (q ≤ 0.05). **e.** Representative fixed-SNV effect profiles for *SMDT1* in TEM cells and *RPS26* in pooled memory T cells. The *SMDT1* profile shows significant deviation from the line connecting the 0 h and 120 h coefficients (gray dashed line). The *RPS26* profile supports a sampled local trough at 40 h relative to 16 h and 120 h. Displayed q values are BY-adjusted gene–population omnibus values for the respective tests. Points indicate reported coefficients; error bars indicate pointwise 95% t confidence intervals without adjustment for representative selection.

Serial sampling revealed additional genetic associations throughout activation. Within 37,173 evaluable gene–population units, cumulative detections increased from 846 before stimulation to 2,320 at 5 d. The number detected at each individual stage peaked at 1,523 at 40 h and declined to 909 at 5 d (**Figure 6b**). The 16 h, 40 h, and 5 d measurements contributed 617, 676, and 181 newly detected associations, respectively, with the largest gains occurring during the first 40 h of activation (**Figure 6c**).

Effect coefficients also changed across activation stages. Direct comparisons with the unstimulated condition identified 15, 12, and four significant gene–population coefficient differences at 16 h, 40 h, and 5 d, respectively. Separate tests detected five absolute-magnitude changes at 16 h and three at 40 h, together with one direction reversal at 16 h (**Figure 6d**). Thus, activation-associated changes encompassed both the strength and direction of genetic regulatory associations.

Individual loci exhibited distinct temporal profiles. In TEM cells, the negative *SMDT1* association weakened markedly at 16 h and became progressively more negative at 40 h and 5 d. This trajectory significantly departed from linear change between the baseline and final measurements (gene–population omnibus BY q = 8.56 × 10⁻¹⁰). In pooled memory T cells, the RPS26 coefficient decreased at 40 h and increased again by 5 d, producing a significant local trough relative to the flanking measurements (gene– population omnibus BY q = 4.64 × 10⁻⁸; **Figure 6e**). These findings show that genetic regulatory associations change in magnitude, direction, and temporal profile during T-cell activation.

## DISCUSSION

### From reference-based to context-resolved variant interpretation

Resolving the functional consequences of somatic mutations in cancer has long been constrained by a dual challenge: the inability of conventional 3’-capture single-cell platforms to broadly recover sparse, endogenous variants across gene bodies, and the reliance of analytical workflows on reference-based, population-frequency heuristics that treat variant impact as a static, cell-agnostic attribute^1,47–49^. Here, Ligo-Seq expands transcript coverage across gene bodies, improving the recovery of endogenous SNVs, while PrismSNV estimates their associated transcriptomic displacement by toggling individual SNV states within a fixed cellular representation. Together, these approaches enable the analysis of patient-private, non-recurrent variants without requiring population-level recurrence or prior clinical annotation. These findings identify the variant-cell pair as a useful unit for interpreting somatic variation, linking variant-associated transcriptomic displacement to cellular identity and regulatory state.

### Private somatic variants exhibit cell-state-dependent molecular effects

By showing that private, non-recurrent somatic SNVs can produce measurable, cell-state-restricted transcriptomic displacement, our results refine the operational binary classification of cancer genomes, which often treats variants lacking detectable positive selection or recurrence as molecularly inert. Our results demonstrate that population-level evolutionary neutrality does not preclude localized, state-dependent transcriptomic displacement at single-cell resolution. It is important to distinguish two complementary dimensions: the driver–passenger framework operates at the level of fitness effects (clonal selection), whereas transcriptomic displacement operates at the level of variant–cell interactions, these are complementary dimensions that need not coincide.

The cell-state-specific transcriptomic displacement we observe is compatible with the nearly neutral theory of molecular evolution^50^, in the sense that molecular effects and population-level fitness consequences need not coincide. Private somatic variants may contribute to intratumoral heterogeneity when they occur in cellular states where the affected gene or regulatory program is active, even if they remain effectively neutral with respect to clonal selection. Notably, a variant may produce substantial displacement in a specific cell state without this automatically conferring a fitness advantage^51^.

### Network buffering as a mechanistic basis for context dependency

The same variant may therefore produce different degrees of displacement depending on the regulatory state of the cell. A likely explanation is that gene-regulatory networks differ in their capacity to buffer local perturbations. When the affected gene or regulatory element belongs to an inactive or redundant module, the transcriptional consequence of an SNV may be minimal^52^. By contrast, the same lesion may become detectable when the cell enters a state in which that module is engaged and contributes to the active transcriptional program. Our trajectory analyses showed that variant-associated transcriptomic displacement varied across cellular states along pseudotime. Differences in the activity of regulatory programs and their capacity to compensate for local perturbations may contribute to this variation. Whether regulatory networks buffer SNV effects at the protein level remains untested. Addressing this question will require platforms that jointly genotype variants, quantify transcriptomic displacement, and assay protein-network activity within the same cell.

### Potential implications for early disease assessment

In the cross-sectional AK-cSCC comparison, premalignant AK lesions showed higher aggregate transcriptomic displacement than matched cSCC lesions (**Figure 4d**). Displacement also varied within cell lineages in AML and MM, with AML HSPCs showing lower scores at later pseudotime positions. These observations suggest that variation in SNV-associated displacement reflects cellular state as well as differences between lesions. In our cohorts, less differentiated or transcriptionally plastic states may be more responsive to somatic variation, whereas established cellular programs may constrain its transcriptomic consequences. Longitudinal or lineage-resolved analyses will be needed to determine whether displacement changes as individual cells move between these states.

### Limitations of the study

Several limitations and opportunities for refinement remain. First, RNA-based SNV detection is constrained by expression level, allelic dropout and incomplete coverage of non-transcribed regions, and conservative filtering may exclude rare but biologically relevant subclonal variants. Paired single-cell DNA and RNA profiling should improve genotype resolution while preserving cell-state information. Second, PrismSNV’s counterfactual estimates assume that SNVs act as independent perturbations conditioned on cellular context. Linked variants and epistatic interactions may violate this assumption; while our LD-like co-occurrence filter (see Methods) addresses statistical redundancy, it does not fully capture biological epistasis. Establishing whether displacement magnitude correlates with clonal fitness advantage requires complementary experimental designs (e.g., lineage tracing with simultaneous transcriptomic profiling). Third, PrismSNV quantifies displacement between decoded transcriptomes after changing an SNV state while holding the inferred cellular context fixed. Because the latent representation is learned from the observed transcriptome, it may already contain expression changes associated with the variant, and the allocation of this information between the latent representation and the SNV branch can influence the estimated displacement. The scores therefore reflect SNV-associated differences within the learned model; their correspondence to transcriptional responses following experimental allele changes requires further validation. Unresolved clonal and cellular-state differences may also influence these estimates. Targeted allele editing with transcriptomic profiling would help assess this correspondence and distinguish gain-of-function from loss-of-function mechanisms^38^.

## STAR★METHODS

### METHOD DETAILS

#### Ligo-Seq chemistry design

Ligo-Seq is a high-throughput single-cell ATAC-RNA co-assay based on scFAST-seq chemistry. The method uses a common cell barcode to label RNA-derived cDNA and accessible chromatin fragments originating from the same nucleus. A single type of barcode hydrogel bead supports both RNA reverse transcription and ATAC-fragment ligation. Each bead releases oligonucleotides containing a cell barcode, a unique molecular identifier (UMI) and a 12 + 7 semi-random primer. The 12-bp fixed sequence functions as an adaptor and downstream PCR anchor, whereas the 7-bp random sequence enables RNA priming without requiring a poly(A) tail.

#### Cell preparation

Fresh tissue specimens were washed in ice-cold PBS (Hyclone, SH30256.01) and dissociated using Collagenase IV (Sigma, C5138-500MG), Dispase II (Roche, 4942078001), Elastinase type I (Sigma, E1250) and trypsin (Sigma, SH30042.01). DNase I (Sigma, 9003-98-9) was added when required according to sample viscosity. After erythrocyte removal (Solarbio, R1010), cell number and viability were assessed using a SeekMate Tinitan Fluorescence Cell Counter (SeekGene, M002C) with AO/PI reagent. Debris and dead cells were removed when required using Miltenyi reagents (130-109-398 and 130-090-101). Fresh cells were washed twice with RPMI 1640 medium (Gibco, 11875119) and resuspended at 1 × 10^6 cells per ml in RPMI 1640 containing 2% FBS (Gibco, 10100147C).

#### Nuclei isolation

Cells were homogenized for 2 min on ice in ice-cold homogenization buffer containing IGEPAL CA-630 (Sigma, I8896), MgCl2 (Macklin, M1028), NaCl (Solarbio, S8210) and RNase inhibitor (Sigma, 3335399001). Homogenates were passed through a 40 *μ*m cell strainer (Sigma, BAH136800040) and centrifuged at 500 × *g* for 5 min at 4 ^∘^C. The nuclear pellet was resuspended in nuclei wash and resuspension buffer containing PBS (Hyclone, SH30256.01) and RNase inhibitor (Sigma, 3335399001). Nuclei were counted using the SeekMate Tinitan Fluorescence Cell Counter with AO/PI reagent. Nuclear integrity and sample purity were examined after staining with 0.4% trypan blue (Sangon Biotech, E607320-0001) under a 40X microscope (Jiangnan Novel Optics, XD-202). Only preparations with intact nuclear envelopes and minimal debris were used for subsequent library construction.

#### Tn5 tagmentation and adaptor design

Single-cell RNA and ATAC libraries were prepared using the SeekOne DD Single Cell Genome Multi-omics (ATAC + RNA) Kit according to the manufacturer’s instructions (SeekGene, K02901). An appropriate number of nuclei was incubated with Tn5 transposase at 37 °C to fragment accessible chromatin and insert adaptor sequences.

Ligo-Seq uses a modified Tn5 adaptor in which PrimerB contains a 5’ phosphorylation modification and an additional 17-bp sequence while retaining a common sequence that can hybridize to the barcode oligonucleotide. This adaptor design enables Tn5-tagged chromatin fragments to receive cell barcodes through DNA ligation during the droplet reaction.

#### Droplet generation and parallel molecular barcoding

After tagmentation, nuclei were mixed with reverse-transcription and DNA ligation reagents and loaded into the sample wells of a SeekOne DD Chip S3. Barcoded Hydrogel Beads and partitioning oil were dispensed into the corresponding wells, and the nuclei, beads and reaction mixture were co-encapsulated using the SeekOne Digital Droplet System. Samples were processed in independent single-sample channels.

The generated emulsions were transferred immediately to PCR tubes and incubated at 25 °C for 30 min followed by 42 °C for 60 min. Within each droplet, bead-derived semi-random primers initiated reverse transcription of RNA molecules to generate barcoded cDNA. In parallel, Tn5-tagged accessible chromatin fragments were ligated to oligonucleotides carrying the same cell barcode and UMI. Thus, RNA-derived cDNA and ATAC-derived DNA from the same nucleus received a shared cellular identity. UMIs in both modalities support molecule-level deduplication; in the ATAC library, the UMI can help distinguish PCR duplicates from independently generated fragments with identical genomic coordinates.

#### Product recovery and pre-amplification

After completion of the droplet reaction, emulsions were broken and the mixed cDNA and ATAC-derived DNA products were recovered and purified using magnetic beads. The recovered molecules were pre-amplified for 7 PCR cycles using a universal forward primer and modality-compatible reverse primers. The purified pre-amplification product was then divided equally for RNA and ATAC library enrichment.

#### ATAC and RNA library construction

For ATAC-library enrichment, one half of the purified pre-amplification product was amplified for 7 PCR cycles using TruSeq Read 1 SeqPrimer and Nextera Read 2 SeqPrimer. Indexed libraries were purified using VAHTS DNA Clean Beads (Vazyme, N411-01) and assessed using a Qubit fluorometer (Thermo Fisher Scientific, Q33226) and a Bio-Fragment Analyzer (Bioptic, Qsep400). For RNA-library enrichment, the remaining half of the purified pre-amplification product was amplified for 8 PCR cycles using TruSeq Read 1 SeqPrimer and template-switching primers. The enriched cDNA was subsequently fragmented, end-repaired, amplified by index PCR and ligated to sequencing adaptors. Indexed libraries were purified using VAHTS DNA Clean Beads and assessed using the Qubit fluorometer and Bio-Fragment Analyzer.

#### Library indexing and sequencing

RNA and ATAC products were converted into independently sequenceable libraries. Dual i5 and i7 sample indexes were incorporated to reduce index hopping during high-throughput sequencing. The cell barcode was positioned in Read 1, avoiding a requirement for a special i5 read orientation or additional customized sequencing cycles.

Both library types were sequenced on the GeneMind SURFSeq 5000 platform using 150-bp paired-end reads.

#### Benchmarking framework

Ligo-Seq was benchmarked against established single-cell technologies on matched mouse brain reference data, independently for the transcriptomic (scRNA-seq) and chromatin-accessibility (scATAC-seq) modalities. Within each modality, every platform was processed through an identical quality-control pipeline and characterised by a panel of per-cell quality metrics. To enable comparison on a common scale, each metric was min-max normalised across platforms (metrics for which lower values indicate higher quality were inverted), and the normalised values were averaged with equal weights to yield a composite quality score in the range 0-1 (scRNA-seq) or 0-100 (scATAC-seq). Metrics that could not be quantified consistently for all platforms were displayed for transparency but withheld from the composite, as detailed below. Analyses were performed in Python 3.10 with scanpy v1.11.5 and anndata v0.10.9, NumPy v1.26.4, SciPy v1.11.4 and scikit-learn v1.3.2; scATAC-seq fragments were processed with SnapATAC2 v2.8.0 and the Ligo-Seq chromatin atlas with ArchR v1.0.3 (peaks called by MACS2 via ArchR); figures were rendered with matplotlib v3.10.8 in Helvetica.

#### Single cell RNA-seq benchmark

Datasets. Nine droplet- and plate-based scRNA-seq technologies were compared: Ligo-Seq, 10x Genomics Chromium v2 and v3, 10x Multiome (gene-expression modality), Smart-seq2, Drop-seq, SPLiT-seq, Microwell-seq and 10x single-nucleus RNA-seq. Publicly available mouse brain datasets were used; full provenance and accession numbers are given in “Code and data availability” below. The latest version of Smart-seq was not included. Count matrices were loaded into *AnnData* objects and, to remove sequencing-depth and cell-number confounds, each was randomly subsampled to a maximum of 20,000 cells (fixed seed). Gene identifiers were harmonised to gene symbols (Ensembl identifiers mapped via the matrix feature metadata; aliasing suffixes removed) and upper-cased for cross-platform matching.

Quality metrics. Four metrics were quantified per platform: (i) sensitivity, the median number of genes detected per cell; (ii) split-half reproducibility, the mean Pearson correlation between pseudobulk profiles of random half-splits of cells over five iterations; (iii) marker silhouette, the silhouette coefficient of marker-defined cell types in principal-component space;(iv) mitochondrial content, the median per-cell fraction of counts mapping to mitochondrial genes (inverted). The marker silhouette is undefined for datasets containing fewer than two marker-resolved cell-type classes; for these platforms (10x v2, 10x v3, Smart-seq2) the silhouette was taken from the directly computed value (0.189, 0.135 and 0.099, respectively) so that all platforms were scored on an identical metric set. Mitochondrial content was computed directly from each count matrix using mitochondrially encoded genes; Smart-seq2 and SPLiT-seq matrices, which do not contain annotated mouse mitochondrial genes, were recorded as not applicable and omitted from this metric rather than assigned a value. The number of cells analysed per sample was additionally reported as a measure of throughput but, as a property of experimental scale rather than of platform chemistry, was not included in the composite.

#### Single cell ATAC-seq benchmark

Datasets and processing. Four scATAC-seq technologies were compared on mouse brain fragment data: Ligo-Seq, TXCI-ATAC (GEO GSM7852211), 10x scATAC (atac_v1_E18_brain_flash_5k) and 10x Multiome (ATAC modality). Fragment files were processed with SnapATAC2 v2.8.0 (ref. Zhang et al., Nat. Methods 2024) against the mouse mm10 assembly with GENCODE vM31 gene annotation. Nuclei were retained at uniform thresholds (larger than 1,000 unique fragments and transcription-start-site enrichment larger than 1 per nucleus), and per-nucleus metrics were computed under identical parameters across platforms.

Quality metrics. Four metrics were quantified per nucleus: transcription-start-site (TSS) enrichment; fraction of reads in peaks (FRiP); number of unique fragments and nucleosome signal. The composite was computed from the three depth-independent library-quality metrics (TSS enrichment, FRiP and nucleosome signal); fragment number, which scales with sequencing depth, was displayed but withheld from the composite, and duplication rate was not used because the TXCI fragments had been de-duplicated upstream. Nucleosome signal was defined as the per-nucleus ratio of mononucleosomal (147-294 bp) to nucleosome-free (<147 bp) fragments. Fragment lengths were computed as the absolute coordinate span, and zero-length insertion records were excluded, to accommodate fragments files in which a fragment is represented in both orientations (as in the merged TXCI file). A pronounced nucleosomal banding pattern is a canonical indicator of successful Tn5 transposition (Buenrostro et al., Nat. Methods 2013; ENCODE ATAC-seq standards); all platforms fell well below the over-transposition exclusion threshold (nucleosome signal >4 in Signac/ArchR), and within this regime Ligo-Seq exhibited the most pronounced mononucleosome peak (mononucleosome fraction 43%, versus 29% for 10x scATAC and 23% for TXCI).

### Patient cohort and sample collection

The study was approved by the Medical Ethics Committee of the First Affiliated Hospital of Xi’an Jiao Tong University (approval No. XJTU1AF2024LSYY-060) and the Second Affiliated Hospital of Xi’an Jiaotong University (approval No. 2024 Ethics Review-Research 004). Written informed consent was obtained from all participants, and all procedures were conducted in accordance with the Declaration of Helsinki.

Skin biopsy specimens were obtained from ten patients with cSCC at the Department of Dermatology, The Second Affiliated Hospital of Xi’an Jiaotong University. AK and cSCC samples were obtained from donors under a protocol approved by the relevant ethics committee. Nine individuals were clinically and histopathologically confirmed to have AK-derived cSCC (progressing from actinic keratosis), and one presented with cSCC arising from a rare dermatological condition. Anatomical sampling sites comprised the face and the dorsum of the hands. For each patient with paired lesions, biopsies were collected from both the AK and the cSCC lesions. Three bone marrow blood samples from three patients with AML and four bone marrow blood samples from four patients with MM were collected from the Department of Hematology, the First Affiliated Hospital of Xi’an Jiaotong University.

In total, 26 discrete samples were processed for sequencing. To resolve coordinated transcriptomic and chromatin-accessibility landscapes, we performed single-cell multi-omics profiling on 16 samples, simultaneously capturing whole-transcriptome scRNA-seq and scATAC-seq profiles from the same individual cells. Ten additional samples were profiled using 3’ single-cell RNA sequencing.

### Cell preparation for single cell 3’ sequencing

After harvesting, tissues were digested with digestive enzymes to prepare and harvest a single-cell suspension. Subsequently, we followed the ‘PIPseq V T10 3’ Single Cell RNA Kit’ user guide and performed the experiment using the ‘Pipseq V T10 3’ Single Cell RNA Kit Bundle’ (PN 20135691, Illumina Inc., San Diego, CA). The next day, we isolated mRNA and reversed transcribed it into cDNA with whole transcriptome amplification. We quantified the concentration of the isolated cDNA using the Qubit HS dsDNA assay kit (Cat. No. Q32851, Thermo Fisher Scientific, Waltham, MA), then diluted it to approximately 1-2 ng/μL for cDNA size quantification using the Agilent 2100 Bioanalyzer (Lot No. 2445). We used the entire volume of cDNA to prepare libraries. After diluting the final libraries at a ratio of 1:10, we quantified their concentrations using the Qubit HS dsDNA assay kit. We then determined the size of the 1:10 diluted libraries using the Agilent 2100 Bioanalyzer with High Sensitivity DNA Reagents. Each library type was sequenced on a different lane of the flow cell using a CNVL00108 NextSeq1000 sequencer.

### Single-cell RNA sequencing data preprocessing

Single-cell transcriptomic data were imported and preprocessed utilizing the Scanpy^53^ framework. During the quality control phase, genes expressed in fewer than 50 cells were filtered out. Cells with fewer than 50 detected genes or a mitochondrial gene fraction exceeding 15% were excluded, and potential doublets were removed from the dataset. Subsequently, highly variable genes (HVGs, top 2000) were identified. Dimensionality reduction was performed using Principal Component Analysis based on the top 15 principal components, followed by unsupervised clustering using the Leiden^54^ algorithm (resolution=0.3). Finally, cluster-specific marker genes were identified by differential expression analysis and used for manual cell-type annotation (**Supplementary File 1**). Copy number variations (CNVs) within the single-cell data were analyzed using the inferCNV (v3.20)^55^ package. Cell-cell communication analysis was performed using CellChat (v2.1.2)^56^, while pseudotime and diffusion pseudotime analyses were conducted using Seurat v5^57^ and Scanpy (v1.11.5)^53^, respectively.

### Single-cell SNV identification and filtering

To identify SNVs from single-cell sequencing data, we implemented a strict variant calling pipeline tailored for transcriptomic data. Raw alignment files (BAM) were first subjected to quality control filtering using SAMtools (v1.23)^58^. We excluded reads that were unmapped, mate-unmapped, secondary alignments, failed quality checks, or marked as PCR duplicates (SAM flag exclusion criteria set to 1804). Additionally, we enforced a minimum mapping quality threshold (*MAPMM* ≥ 30) to minimize artifacts arising from ambiguous alignments^59^.

Pileup files were generated from the filtered BAMs using samtools *mpileup* with base alignment quality (BAQ) computation disabled to preserve original base quality scores. SNVs were subsequently called using VarScan2^60^ in *mpileup2snp* mode. To classify a locus i as a putative variant, we required the sequencing data to satisfy the following criteria:

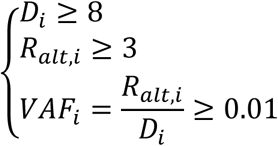

where *D_i_* denotes the total read depth at locus *i*, *R_alt_*_,*i*_ represents the number of reads supporting the variant allele, and *VAF_i_* is the variant allele frequency. To distinguish true genomic variants from post-transcriptional modifications, the resulting VCF files were cross-referenced with a database of known RNA editing sites using BEDTools (v2.31)^61^, and overlapping variants were subsequently excluded. Next, variant annotation was performed using ANNOVAR (v2020-06-07)^62^ against the gnomAD^19^ database (v4.1, hg38). To prioritize rare variants, all SNVs with a population minor allele frequency (MAF) exceeding 0.001 were discarded.

To establish an empirical allele frequency (AF) threshold for filtering putative germline variants from RNA-derived SNVs, we utilized germline SNVs identified via whole-exome sequencing (WES) as an orthogonal reference standard. RNA-derived SNVs were rigorously cross-referenced with the WES-defined germline SNVs, requiring exact concordance of chromosome, genomic coordinate, reference allele, and alternative allele to generate a high-confidence intersected SNV set.

Subsequently, we extracted the RNA-derived AFs within this concordant set and defined our germline filtering threshold, *τ*, as the 0.5th percentile of their empirical distribution:

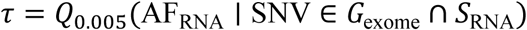

Here, *G*_exome_ and *S*_RNA_ denote the sets of WES-derived germline SNVs and RNA-derived SNVs, respectively. By this formulation, approximately 99.5% of WES- supported germline RNA SNVs exhibit an AF exceeding *τ*, establishing this threshold as a highly conservative lower bound for germline variant AFs within the RNA callset. Consequently, in all downstream analyses, RNA-derived SNVs with AF ≥ *τ* were classified as germline variants and systematically excluded.

Finally, to mitigate false positives arising from sequence homology and somatic hypermutation, variants mapped to immunoglobulin and T-cell receptor V(D)J recombination loci were also removed from the final dataset.

### Single-cell genotyping and matrix construction

To systematically map identified variants to individual cells, we first constructed a unified, non-redundant catalogue of target SNVs. Variant calls from all samples were filtered to retain only those with an allele frequency below a specified threshold (e.g., user-defined parameter *P_freq_*) to prioritize rare or sub-clonal somatic mutations, and then merged into a master list.

We then performed cell-specific genotyping by interrogating the aligned reads at these target loci using the pysam library. Reads were assigned to cells based on their corrected cell barcode (CB) tags. To ensure high-confidence genotyping, we excluded reads with base quality scores less than 20 at the query position and ignored loci with excessively high pileup depth (> 50,000) to avoid alignment artifacts in repetitive regions.

For each cell *i* and variant *jj*, we constructed a genotype matrix *G* using a ternary encoding scheme to represent the allelic status:

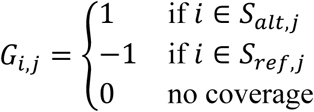

where *S_alt_*_,*jj*_denotes the set of barcodes supported by at least one read carrying the alternative allele, and *S_ref_*_,*jj*_denotes the set of barcodes supported by reads carrying the reference allele.

To quantify the reliability of entries designated as REF-only (−1) in the cell-by-variant matrix, we computed the posterior probability that a given cell truly carries the reference allele, conditioned on the observed read evidence. Let *a_ijj_* and *r_ijj_* denote the number of ALT- and REF-supporting observations, respectively, for cell *i* at single nucleotide variant (SNV) site *jj* . Observations were defined as unique molecular identifier (UMI) counts where available, or raw read counts otherwise. An entry was initially assigned a REF-only state (−1) if *a_ijj_* = 0 and *r_ijj_* ≥ 1.

We modeled the ALT observations using a binomial likelihood under two latent states: true reference (*Z_ijj_* = REF) and true variant carrier (*Z_ijj_* = 1). Letting *n_ijj_* = *a_ijj_* + *r_ijj_*, the conditional distributions are:

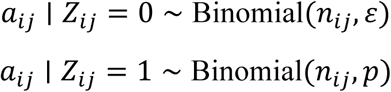

where *εε* is the effective ALT error rate for a true reference allele, and *p* is the expected ALT fraction for a true carrier (e.g., *p* ≈ 0.5 for heterozygous loci lacking allelic bias).

The cell-level prior probability of being a carrier, *π_jj_* = Pr(*Z_ijj_* = 1), was informed by the sample-level variant allele frequency (VAF*_jj_*) derived from the bulk VCF. We applied a scaled and capped mapping:

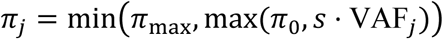

where *π*_0_ serves as the default prior for sites without a defined sample-level VAF, *s* is a scaling factor (typically *s* ≈ 2 under a heterozygous approximation), and *π*_max_ imposes an upper bound on the prior.

For REF-only entries (*a_ijj_* = 0, *r_ijj_* ≥ 1), the posterior probability of the true reference state is thus:

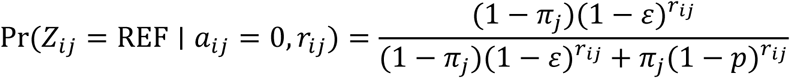

This posterior probability was utilized as a continuous reliability metric for all −1 calls, allowing for the evaluation of its distribution, via histograms and quantiles, across the dataset. To enforce stringent quality control, low-confidence REF-only entries could optionally be reassigned to an unknown state (0) by applying a user-defined threshold to this posterior probability. The resulting matrices from all samples were concatenated into a unified *AnnData* object for downstream analysis.

### Unsupervised representation learning and batch effect correction

To derive robust, low-dimensional cellular representations invariant to technical covariates, we implemented a deep generative framework based on a Variational Autoencoder (VAE) architecture^63^. This pre-training stage serves to capture the intrinsic biological manifold of the single-cell transcriptomic data while reducing batch-associated variation in the latent representation.

The RNA backbone was an MLP-based variational autoencoder. The encoder comprised Linear–LayerNorm–ReLU–Dropout–Linear–LayerNorm–ReLU, followed by separate linear heads for the posterior mean and log variance in a 128-dimensional latent space. The latent representation was sampled as

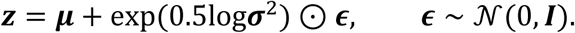

To account for the overdispersion and sparsity inherent in scRNA-seq UMI counts, we modeled the reconstruction loss using a Negative Binomial (NB) likelihood. The decoder maps the latent representation **zz**, conditioned on batch covariates, back to the parameter space of the gene expression distribution. The decoder produced the gene-wise mean expression, whereas dispersion was represented by a learned gene-specific parameter shared across cells. The reconstruction objective is defined as the negative log-likelihood of the NB distribution:

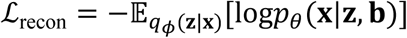

where **bb** represents the batch indices. The dispersion **θ** is constrained to be positive using a softplus activation to ensure numerical stability.

Dual-Strategy Batch Correction

We employed a two-pronged approach to integrate data from diverse sources and reduce batch-associated variation:

1. Conditional Decoding: Categorical batch covariates are projected into a learnable embedding space (dimension *d* = 16). These batch embeddings are concatenated with the latent vector **zz** to form the input for the decoder. This conditional VAE formulation encourages the model to use **zz** to encode biological variation while relying on the batch embedding to reconstruct technical artifacts.
2. Adversarial Domain Adaptation: To further reduce batch-related information from the latent space, we introduced an adversarial training scheme via a Gradient Reversal Layer (GRL). A discriminator network functions to predict the batch identity solely from the latent representation **zz** . During backpropagation, the gradients from the discriminator are reversed (multiplied by −*λ*_adv_) before passing to the encoder. This minimax game enforces the generation of a batch-invariant latent distribution *q*(**zz**) that retains biological signal but confuses the batch discriminator.

### Optimization and Training

The pretraining objective combined negative binomial reconstruction, KL regularization and batch-classification cross-entropy:

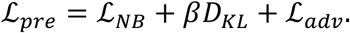

The negative binomial likelihood accommodates the overdispersion of single-cell count data, while KL regularization constrains the posterior distribution and promotes a smooth latent representation. Batch-classification loss was applied through a gradient-reversal layer: the discriminator learned to identify batch labels, whereas the adversarial gradient propagated to the encoder was multiplied by −*λ*_adv_. Together with the batch-conditioned decoder, this design allowed measured batch information to support reconstruction while discouraging the latent representation from retaining readily batch-predictive features. The resulting latent space therefore provided a more stable cellular background for downstream SNV-conditioned modeling.

The KL weight *β* and the GRL coefficient *λ*_adv_ were increased linearly during warm-up. Gradually introducing these constraints prevented them from dominating reconstruction during the early stages of training and reduced instability before the encoder had learned the major expression structure. The optimizer used a linear learning-rate warm-up followed by cosine annealing to support stable initial updates and gradual convergence.

### SNV perturbation effect modeling

Building on the pretrained RNA backbone, the SNV-conditioned model initialized all shape-matched encoder and decoder parameters from the backbone checkpoint and retained the same negative binomial observation model. To preserve the separation established during RNA-only pretraining, the encoder was frozen throughout SNV training, preventing its parameters from adapting to the observed SNV assignments. The decoder, batch embedding and SNV-specific modules remained trainable; however, the decoder was optimized at a learning rate of 10⁻⁴, one order of magnitude lower than the 10⁻³ learning rate used for the remaining trainable parameters. This asymmetric optimization was designed to preserve a stable RNA-derived cellular context while allowing the SNV branch to learn an incremental, context-dependent correction, rather than allowing joint optimization to redistribute the complete expression signal arbitrarily between the cellular latent state and the SNV-specific pathway. Each SNV *v* was represented by a learnable embedding *e_v_* . For each covered cell–SNV pair (*c*, *v*), we computed a context-dependent representation:

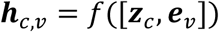

and an independent sigmoid gate:

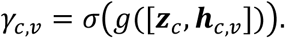

Unlike softmax attention, these gates were not normalized across SNVs within a cell. The signed SNV contribution was

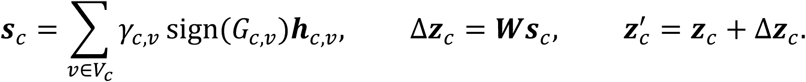

The projection W contained no bias, ensuring that zero gated contribution produced no latent offset. This exact no-op property prevented the SNV pathway from introducing a systematic latent shift in the absence of gated SNV evidence and preserved *z_c_* as the unperturbed model state. To scale to large variant sets, training and inference processed only non-zero cell-SNV pairs in chunks and accumulated their signed, gated embeddings back to the corresponding cells. This avoided dense cell-by-SNV-by-latent tensors and required no per-cell softmax or log-sum-exp normalization.

### Latent-contribution-based SNV prioritization

For each covered cell–SNV pair, we quantified the latent displacement produced by flipping the focal allelic state as

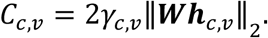

Using the encoder posterior mean as the cellular latent state, the latent-contribution score of SNV *v* was defined as

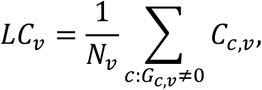

where *N_v_* is the number of covered cells. SNVs were ranked by *LC_v_* globally and within each cell type, and the top-ranked variants were retained for decoder-based counterfactual scoring. Cell-type-specific screening required more than 20 cells in each allelic state. The latent-contribution score was used only for preselection and was distinct from the final transcriptomic displacement score.

### Counterfactual SNV effect scoring

For each selected SNV *v* and covered cell *c*, we held the encoder posterior mean, batch covariates and all other SNV states fixed. Importantly, the same cellular latent state *z_c_* was used for both decoding operations, so the modeled cell identity and cellular context remained constant throughout the paired comparison. The expression profile was decoded first using the observed allelic state and then after flipping only the focal state, *G_c_*_,*v*_ ← −*G_c_*_,*v*_, yielding 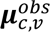 and 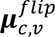. We defined model-derived transcriptomic displacement as the difference between the two reconstructed expression profiles obtained by flipping the focal SNV state while holding the inferred cellular context, batch covariates and all other SNV states fixed. Its magnitude and pattern were quantified using Euclidean and cosine distances, respectively:

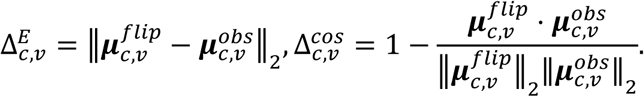

Euclidean and cosine distances were reported as complementary measures of SNV- state-associated transcriptomic displacement. Euclidean distance quantifies the overall magnitude of change between the two decoded expression profiles and is therefore sensitive to absolute differences across genes. Cosine distance quantifies their angular dissimilarity, emphasizing changes in the relative multigene expression pattern while being less dependent on overall vector magnitude. Scores were retained for individual covered cell-SNV pairs and summarized across covered cells for SNV- or cell-type-level analyses. Used in combination, these complementary metrics quantify the relative magnitude and pattern of SNV-associated transcriptomic perturbation within the learned cellular context.

### Cross-method evaluation using a common RNA distance-delta metric

To compare different SNV-prioritization strategies on a common scale, candidate SNVs were first restricted to a shared coverage-eligible universe containing at least 30 ALT-supported and 30 REF-supported cells across the aligned dataset. Each method then prioritized candidates according to its own predefined criterion. PrismSNV candidates were defined as the top 10% of SNVs ranked by decoder-based counterfactual perturbation score within each cell type. VEP and SnpEff candidates were selected using their respective HIGH-impact classifications, whereas MutationTaster candidates were selected using its Deleterious classification. These method-specific scores and annotations were used only for candidate selection; the RNA distance-delta metric described below was not used to define any candidate set.

All method-prioritized candidate sets were subsequently evaluated using the same RNA-based endpoint. For each evaluable SNV *v* in cell type *t*, we calculated the Euclidean distance between the mean observed RNA-expression vectors of ALT- supported and REF-supported cells:

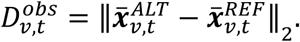

To estimate the distance expected from random separation of the same numbers of cells, ALT and REF group labels were randomly reassigned within the same cell type while preserving the observed group sizes. The RNA distance delta was defined as:

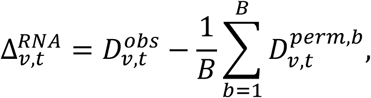

where 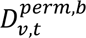 denotes the Euclidean distance obtained in randomization *b*. Positive values therefore indicate that the observed ALT–REF transcriptomic separation exceeds that expected from random cell-group assignments. For each method and cell type, RNA distance deltas were averaged across the corresponding method-prioritized SNVs. As an additional background reference, 100 eligible SNVs were randomly sampled from the same candidate universe in each of 100 repetitions. The same RNA representation, distance calculation and random-background procedure were applied to all four candidate sets, providing a common evaluation endpoint independent of their method-specific prioritization criteria.

### SNV-matched reconstruction regularization

During SNV training, the matched RNA–SNV profile was used for the primary negative binomial reconstruction and KL objectives. To constrain the SNV branch to act as a residual update rather than an alternative encoding of the complete transcriptome, we imposed a soft upper bound on the relative perturbation magnitude:

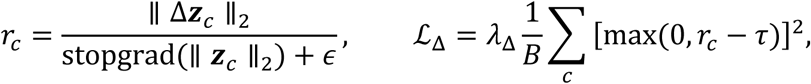

with *τ* = 0.8 and *λ*_Δ_ = 100.

This penalty remained inactive for perturbations below the specified relative threshold and was applied only when the SNV-induced update became disproportionately large relative to the cellular latent state. It therefore preserved context-dependent SNV responses while limiting solutions in which the SNV pathway overwhelmed or replaced the pretrained representation of cellular identity. We additionally used a same-batch derangement loss to prevent the model from achieving comparable reconstruction while ignoring the supplied SNV assignment. For each RNA profile, the matched and mismatched SNV configurations were evaluated using the same encoder posterior mean and batch context. Because these paired alternatives differed only in the supplied SNV configuration, a model relying exclusively on the RNA-derived latent state would obtain no ranking advantage. Satisfying the ranking margin therefore required the explicit SNV branch to retain assignment-specific information. Restricting negative configurations to other cells from the same batch further reduced the possibility that the ranking objective could be satisfied through batch-associated differences:

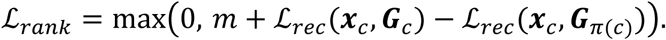

Pairs differing at fewer than five SNV entries were excluded. The ranking term was introduced after 10 epochs, with its margin and weight linearly increased to 0.05 over the following 10 epochs. The final SNV-training objective was:

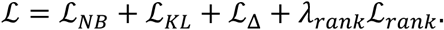

Encoder freezing, limited decoder adaptation and residual regularization constrain the SNV branch to learn an incremental reconstruction update over the RNA-derived cellular context. The matched-versus-shuffled objective specifically trains this update to retain information from the observed SNV configuration, rather than allowing reconstruction to rely on the RNA representation alone. Subsequent single-SNV toggling measures the reconstruction change attributed by the model to the focal variant while keeping the cellular context and other SNV states fixed.

### LD-like SNV co-occurrence filtering

To reduce redundancy among highly co-occurring SNVs, we applied a post-processing filter to the cell-type–stratified perturbation score matrix. Within each cell type, duplicated (cell type, SNV) entries were resolved by retaining the instance with the highest Euclidean perturbation score. SNV calls were aligned to RNA-based cell-type annotations by shared barcodes, and only ALT-supported cells (value = 1) were considered carriers.

For each cell type and sample, co-occurrence between SNV pairs was quantified using carrier-cell sets. We computed the Jaccard index:

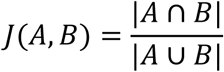

and the overlap coefficient:

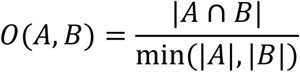

where *A* and *B* denote the carrier sets of the two SNVs. Pairs with ≥20 carriers per SNV, ≥20 shared carriers, *J* ≥ 0.85, and *O* ≥ 0.95 were considered redundant and represented as edges in a graph. SNVs were ranked by perturbation score, and lower-scoring SNVs directly connected to higher-scoring ones were removed.

This filter captures LD-like redundancy based on single-cell co-occurrence patterns rather than haplotype-based linkage disequilibrium.

### Validation of counterfactual inference and cell-type specificity

To validate the cell-type-specific effects of SNVs, we quantified counterfactual perturbation magnitude using the Euclidean distance between the decoded expression profiles under the reference and alternative allelic states, defined as:

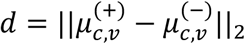

All SNVs passing the prespecified technical filters were included in the cell-type-specificity analysis; no preselection based on the final counterfactual displacement score was performed. For each SNV, cells with ALT support (*G_c_*_,*v*_ = +1) were first used to identify the patient and sample blocks in which the variant was observed. Subsequent scoring was restricted to cells from these host blocks with either ALT or REF evidence (*G_c_*_,*v*_ ≠ 0), thereby avoiding the evaluation of patient-private variants in unrelated patients containing only hypothetical reference backgrounds. Within each eligible cell type, up to 100 covered cells were randomly sampled, and cell types represented by fewer than 3 covered cells were excluded.

Cell-type dependence was quantified using eta-squared and the Kruskal–Wallis statistic. To construct an exchangeability-preserving null distribution, cell-type labels were permuted among covered cells within each host sample block, while RNA profiles, SNV states, batch labels and counterfactual distances remained fixed. The counterfactual distance for each cell–SNV pair was calculated once and reused across all permutations. For each SNV and statistic *T*, the empirical P value was calculated as 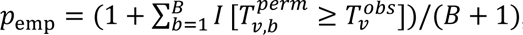, where *B* = 1000 was the number of blocked permutations. Empirical P values based on eta-squared were corrected across all tested SNVs using the Benjamini–Hochberg procedure, and empirical q < 0.05 was used as the primary definition of cell-type specificity. Nominal Kruskal–Wallis P values were retained as descriptive statistics only. Patient-private SNVs were subsequently summarized within each host patient, and cohort-level prevalence was reported across patients rather than requiring recurrence of the same SNV locus.

For each SNV, relative displacement amplitude was calculated as:

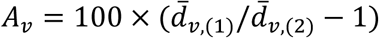

where *d̄*_*v*, (1)_ and *d̄*_*v*, (2)_ are the two largest cell-type mean counterfactual distances.

SNVs exceeding the 10% or 30% thresholds were required to have an empirical BH- adjusted *η*^2^ *q* < 0.05.

### Single-cell ATAC-seq data processing and peak-gene linkage analysis

For each scATAC-seq sample, the paired RNA expression data from previous step and ATAC peak count matrices were integrated by harmonizing cell barcodes across modalities. Subsequently, a signac^64^ ChromatinAssay object was constructed for each sample, incorporating the peak count matrix, fragment alignments. The individual datasets were then merged into a unified dataset. Peaks detected in fewer than 50 cells or those mapped to non-standard contigs were excluded to ensure robust downstream analysis. Genomic metrics, including peak GC content and sequence length, were calculated based on the GRCh38 reference. Following filtering, the ATAC-seq data underwent term frequency-inverse document frequency (TF-IDF) normalization, followed by dimensionality reduction via singular value decomposition (SVD) to generate latent semantic indexing (LSI) components. Previously established cell type annotations were then mapped to the integrated multi-omic dataset.

To identify putative regulatory interactions, peak-gene linkages were inferred using the LinkPeaks algorithm. The search space was restricted to a 500-kilobase (kb) window surrounding the transcription start site (TSS) of each target gene. Significant peak-gene associations were evaluated either globally across all cells or within specific cell populations, defined by a correlation P-value threshold of < 0.05. Finally, joint genomic tracks visualizing the chromatin accessibility landscape, aligned gene expression profiles, and significant peak-gene linkages were generated using the CoveragePlot function, encompassing a 100 kb flanking region upstream and downstream of the target loci.

### Motif annotation and SNV disruption analysis

To establish a chromatin accessibility landscape, de novo peak calling was performed across all integrated cells utilizing MACS2^65^. The resulting unified peak set was subsequently quantified into a peak-cell matrix. To identify potential regulatory elements within these accessible regions, motif annotations were incorporated using vertebrate transcription factor (TF) position frequency matrices (PFMs) obtained from the JASPAR2020^66^ database. The top 30% of high perturbation SNVs identified by PrismSNV within each cell type were retained for downstream analysis. Following the standardization of SNV coordinates to the human reference genome (GRCh38), genomic intersections were performed to map these variants to both the robust ATAC- seq peaks and the predicted TF binding sites.

The potential of these mapped SNVs to disrupt TF binding was systematically assessed using the motifbreakR^67^. Disruption scores were evaluated by comparing the binding affinities of the reference and alternate alleles against a curated motif database, applying an information content P-value threshold of <1e-4. Variants exhibiting profound predicted effects—characterized by a significant gain or loss of motif binding affinity— were identified to illustrate sequence-level regulatory disruptions. Finally, the cell-type-specific distributions of perturbation scores and the overall disruption rankings of TF motifs were quantified to evaluate the impact of these variants on the cis-regulatory network.

### AlphaGenome-based prediction of SNV regulatory effects

To estimate the regulatory impact of candidate SNVs, we queried AlphaGenome using the Python SDK. SNVs were first extracted from the PrismSNV cell-type perturbation table, deduplicated, and reformatted as genomic variants with chromosome, position, reference allele, and alternative allele. Variants located outside primary human chromosomes were excluded. For each remaining SNV, AlphaGenome predictions were generated with the human model using scRNA-seq and scATAC-seq scorers. scRNA-seq effects were predicted using a 1-Mb sequence context centered on each SNV, whereas ATAC-seq effects were predicted using a 16-kb context. We retained skin-relevant tracks by matching tissue annotations related to skin, suprapubic skin, lower-leg skin, epidermis, dermis, keratinocytes, melanocytes, or foreskin. For each SNV, AlphaGenome effect scores were summarized across retained skin-related tracks using the absolute quantile score, including mean, median, interquartile range, and maximum values. These summaries were used as sequence-based estimates of SNV regulatory effect and compared with PrismSNV-derived single-cell perturbation scores.

### Generation of stable COL3A1 knockdown fibroblasts

Three short hairpin RNAs (shRNAs) targeting the 3’ untranslated region (3’UTR) of human COL3A1 were designed with the following target sequences:

- shRNA1, 5’-ATGTCTCAATGGTGCTATAAT-3’
- shRNA2, 5’-CTCCTGGAAGATTGGTTTAAA-3’
- shRNA3, 5’-ACTTGCATTCAGAACATAAAT-3’

The three shRNAs were incorporated into the pGC-CMV-BSD-H1-COL3A1-shRNA1/2/3 lentiviral construct. A corresponding non-targeting shRNA construct was used as the negative control. Recombinant constructs were verified by sequencing before lentiviral production.

Lentiviruses were generated by co-transfecting HEK293T cells with the shRNA expression construct and the packaging plasmids psPAX2 and pMD2.G. Culture medium was replaced 12 h after transfection, and viral supernatants were collected at 48 and 72 h and concentrated by ultrafiltration. Human skin fibroblasts (HSFs) were maintained in DMEM supplemented with 20% fetal bovine serum and 1% penicillin– streptomycin. Cells were seeded in six-well plates and transduced at approximately 30– 50% confluence in the presence of 5 μg/mL polybrene. Medium was replaced after 24 h. At 72 h after transduction, cells were passaged and selected with blasticidin (4 μg/mL) for 4 days, with untransduced cells included to monitor selection efficiency. Surviving populations were expanded for subsequent experiments. COL3A1 knockdown was assessed by quantitative PCR using the 2^−ΔΔCt^ method with β-actin as the reference gene and by western blotting. The COL3A1 quantitative PCR primers were:

- 5’-CGAAGCCGAAGACCTGTTCTA-3’ (forward)
- 5’-GGGCAAGTGTGGACTGTTCC-3’ (reverse).

### Construction and stable expression of WT and mutant COL3A1

The human COL3A1 coding sequence was amplified from a COL3A1 expression plasmid and inserted into a pSBbi-derived Sleeping Beauty vector carrying a C-terminal 3×FLAG tag and a puromycin resistance cassette. The p.G726E and p.G858D substitutions were introduced using mutation-containing primers, followed by seamless assembly of the amplified fragments into the same vector backbone. WT and mutant constructs contained the COL3A1 coding sequence without the endogenous 3’UTR targeted by the knockdown shRNAs. Recombinant clones were verified by sequencing, and purified plasmids were prepared for cell transfection.

Established COL3A1-knockdown and non-targeting control HSFs were used as recipient cells. For each COL3A1 expression condition, the corresponding WT or mutant expression plasmid (2 μg) was co-transfected with the SB100X transposase plasmid (0.2 μg) using a lipid-based transfection reagent. Culture medium was replaced 12 h after transfection and refreshed at 48 h. Cells were passaged at a 1:3 ratio at 72 h and selected with puromycin (2 μg/mL) for 3 days. Selected cell populations were expanded, and COL3A1 expression was assessed by quantitative PCR and western blotting using a FLAG-tag antibody before downstream phenotypic analyses.

### Western blot analysis

HSFs were collected in ice-cold PBS and lysed in RIPA buffer supplemented with protease inhibitors and PMSF on ice for 10 min. Lysates were clarified by centrifugation at 13,000 rpm for 15 min at 4°C, and protein concentrations were determined using a BCA assay. Equal amounts of protein (20 μg per lane) were separated by SDS–PAGE and transferred onto PVDF membranes. Membranes were blocked with 5% skim milk in TBST for 1 h at room temperature and incubated with an anti-COL3A1 primary antibody (1:1,000) overnight at 4°C. After three washes with TBST, membranes were incubated with an HRP-conjugated goat anti-rabbit IgG secondary antibody (1:5,000) for 1 h at room temperature. Membranes were washed again, and protein bands were visualized using enhanced chemiluminescence.

### Immunofluorescence staining

HSFs were cultured on coverslips, washed with PBS and fixed with 4% paraformaldehyde for 15 min. Cells were permeabilized with 0.5% Triton X-100 in PBS for 20 min at room temperature and blocked with 1% bovine serum albumin in PBS for 30 min. Coverslips were incubated with an anti-Collagen III (COL3A1) antibody (Abcam, ab7778, rabbit polyclonal, 1:100) overnight at 4°C, washed three times with PBST, and subsequently incubated with FITC-conjugated goat anti-rabbit IgG (H+L) (Boster, BA1105, 1:500) for 1 h at room temperature in the dark. After washing, coverslips were mounted with mounting medium and examined by fluorescence microscopy.

### CCK-8 assay

CCK-8 assays were performed on HSFs carrying COL3A1 knockdown alone, knockdown plus empty vector, or knockdown plus WT, G726E or G858D COL3A1. Cells carrying non-targeting shRNA and empty vector were included as a control. Cells were cultured in assay plates and assessed at 24, 48 and 72 h. At each time point, CCK-8 reagent was added at a volume equal to 10% of the culture medium, followed by incubation for 2 h. Absorbance was measured at 450 nm. Wells containing culture medium and CCK-8 reagent without cells served as blanks, and their absorbance was subtracted from the corresponding sample readings. Results were expressed as blank-corrected OD450 values.

### Analysis of dynamic SNV-gene eQTL effects during T-cell activation

Published CD4⁺ T cell cis-eQTL summary statistics were obtained from Zenodo (DOI: 10.5281/zenodo.6006796). Analyses included TN, TCM, TEM, and TEMRA cells at 0 h, 16 h, 40 h, and 5 d, with pooled naive and memory populations analyzed separately. Single-nucleotide substitutions were matched across contexts using gene and variant identifiers. Reported coefficients (β), standard errors (SE), and nominal P values were retained.

Coefficients were compared across all time-point pairs within each population and across population pairs at matched times. For two estimates, we calculated *R* = |*β*_1_ − *β*_2_|/(*SE*_1_ + *SE*_2_). P values were obtained by summing the corresponding two-sided Student’s t-tail probabilities and capping the result at one, using context-specific residual degrees of freedom inferred from the published statistics. Within each gene, Bonferroni correction accounted for all eligible SNV-by-comparison tests, followed by Benjamini–Yekutieli (BY) correction across genes at q ≤ 0.025 per comparison family. One fixed representative SNV per identified gene was displayed using signed coefficients on a shared color scale.

Eligible SNV–gene pairs were intersected across all four time points within each population. Association detection used within-gene cis-SNV Bonferroni correction followed by BY correction across gene–population–time tests (q ≤ 0.05). Current, cumulative, and first-detection counts were summarized from these results. Each post-stimulation condition was also compared with 0 h for signed coefficient differences, absolute-magnitude differences, and direction reversals. Magnitude tests applied the same t-tail procedure to ||*β_t_*| − |*β*_0_||/(*SE_t_* + *SE*_0_); reversals were assessed using directional intersection–union tests. Each endpoint underwent cis-SNV Bonferroni correction followed by BY correction across gene–population–comparison events at q ≤ 0.05.

Linear contrasts tested deviations of the 16 h and 40 h coefficients from the line connecting the 0 h and 120 h endpoints (Figure 6e). Local extrema were assessed by intersection–union tests comparing the middle coefficient with both neighbors in the (0, 16, 40 h) and (16, 40, 120 h) triplets. Gene–population omnibus tests incorporated all eligible SNVs and temporal contrasts through Bonferroni correction, followed by BY correction at q ≤ 0.05 separately for each endpoint and population family. Representative profiles retained the same SNV across time, with error bars indicating pointwise 95% t confidence intervals.

### Cross-platform perturbation consistency analysis

To evaluate the reproducibility of PrismSNV-derived perturbation scores between 3’ scRNA-seq and full-length scRNA-seq platforms, clusters from each dataset were annotated to shared major cell types using curated marker-gene panels. Clusters with insufficient marker support were annotated as unassigned or low-information and excluded from cross-platform comparisons.

For cell-level comparisons, perturbation scores were summarized within each platform by calculating the median score across cells belonging to the same major cell type. Cell types were then ranked according to their median perturbation scores, and cross-platform consistency was assessed by comparing cell-type-level score distributions and rank concordance between platforms. For gene-level comparisons, SNV perturbation scores were collapsed to genes by calculating the median perturbation score of SNVs located within each gene, separately for each platform and major cell type. Genes were then ranked by their aggregated perturbation scores, and cross-platform agreement was evaluated using rank correlation and overlap of top-ranked perturbation-associated genes.

For functional comparisons, top-ranked perturbation-associated genes from selected major cell types were used for pathway enrichment analysis with g:Profiler^68^ against Gene Ontology Biological Process, Reactome and WikiPathways databases. Enriched terms were compared between platforms using shared-term overlap and concordance of enrichment significance. Related enriched terms were manually grouped into broader biological themes, including epidermal differentiation, extracellular matrix organization, angiogenesis, cell adhesion and immune crosstalk.

### Computational hardware and run time

All computational experiments were conducted on a high-performance computing workstation. The hardware infrastructure comprises dual Intel Xeon Silver 4316 processors (40 physical cores, 80 logical threads, 2.30 GHz base frequency) and 512 GB of DDR4 RAM. For parallel processing and acceleration, the system is equipped with a quad-configuration of NVIDIA GeForce RTX 4090 GPUs, offering a cumulative 96 GB of GDDR6X memory. The software environment operates on a Linux-based platform, utilizing NVIDIA driver version 580.76.05 and the CUDA 12.4 toolkit. It took approximately 5 hours on this workstation to evaluate the perturbation effects of 10,000 candidate SNV sites across 99,825 cells over 300 epochs, with 14,313 genes per cell.

## RESOURCE AVAILABILITY

### Lead contact

Requests for further information and resources should be directed to and will be fulfilled by the lead contact, Kai Ye.

### Materials availability

All samples used in this study were obtained through patient donation following approval by the institutional ethics committee and after obtaining informed consent from the patients.

### Data and code availability

The code for PrismSNV is publicly available at https://github.com/xjtu-omics/PrismSNV, and the related documentation can be openly accessed at https://prismsnvdoc.readthedocs.io/en/latest/. The code for benchmark is publicly available at https://github.com/xjtu-omics/scRNA_ATAC_analysis. All single-cell ATAC-seq data and single-cell full-length transcriptome sequencing data related to AK-cSCC, MM and AML in this study have been deposited in the National Genomics Data Center database (BioProject accession: PRJCA060066; GSA-Human accession numbers: HRA021777 and HRA017520). Single-cell sequencing data for cSCC can be obtained from the Gene Expression Omnibus (GEO) database (GSE193304 and GSE200334), and single-cell sequencing data for head and neck squamous cell carcinoma can be obtained from GEO database (GSE286827). Additional 3’ single-cell RNA-seq datasets used for cross-cancer evaluation were also obtained from GEO, including intrahepatic cholangiocarcinoma (GSE138709), bladder cancer (GSE135337) and breast cancer (GSE195861). The raw scRNA-seq data for the T-cell dataset are available from the European Genome-phenome Archive (accession number EGAD00001008197). Genotype data have been deposited in the European Genome-phenome Archive (accession number EGAD00010002291). Processed data are available at https://trynkalab.sanger.ac.uk.

## ACKNOWLEDGMENTS

K.Y. is supported by the National Key R&D Program of China (grant no. 2022YFC3400300) and the National Natural Science Foundation of China (grant nos. 32125009 and 32430017). S. G. is supported by the Shaanxi Provincial Innovation Team (No. 2022TD-48) and the Key Industrial Innovation Chain of Shaanxi Province (No. 2024SF-ZDCY L-01-14). Peng Jia is supported by the National Natural Science Foundation of China (grant No. 32400509), the China Postdoctoral Science Foundation (grant nos. 2025M772627, GZB20250610), the National Key R&D Program of China (2025YFC3410300). X.Y. is supported by the National Natural Science Foundation of China (grant no. 32422019), the Natural Science Foundation of Shaanxi Province (grant no. 2024JC-JCQN-28), and the Fundamental Research Funds for the Central Universities (xzy012024088), the Scientific Research Program of Shaanxi Provincial Department of Education (23JK0290). P. Z. is supported by the National Natural Science Foundation of China (grant no. 325B2021). D.M. is supported by the China NSFC projects under contract 62476214 and Tianyuan Fund for Mathematics of the National Natural Science Foundation of China (Grant No. 12426105).

## AUTHOR CONTRIBUTIONS

K.Y. supervised the research; P.S. developed the algorithm and software; Y.G., J.Y. and G.S. developed Ligo-Seq; C.L. and X.X.L. benchmarked Ligo-Seq; D.M., P.S., P.J., H.L., and Q.X. contributed to the assessment and analysis of the deep-learning model; J.Z.L. and Z.S.Y. evaluated the validity of the computational model; S.G., M.L., and X.N.L. provided clinical diagnoses and collected samples; X.J.L. processed the clinical specimens, prepared single-cell suspensions, and performed quality control; R.Z., P.S., and D.X. designed the cellular experiments; K.Y., S.G., P.J., D.M., and X.Y. provided constructive suggestions for the model analysis; Y.G. supported scRNA-seq data preprocessing; P.S., X.X.L., P.J., S.C., T.X., P.Z., Z.Z., S.W., Y.W., M. L. and Y.Z. contributed to downstream data analysis and visualization; K.Y. and P.S. wrote the paper with input from all other authors; S.J.B. and X.Y. reviewed and edited the manuscript. All authors read and approved the final manuscript.

---

**Supplementary Figure 1.**
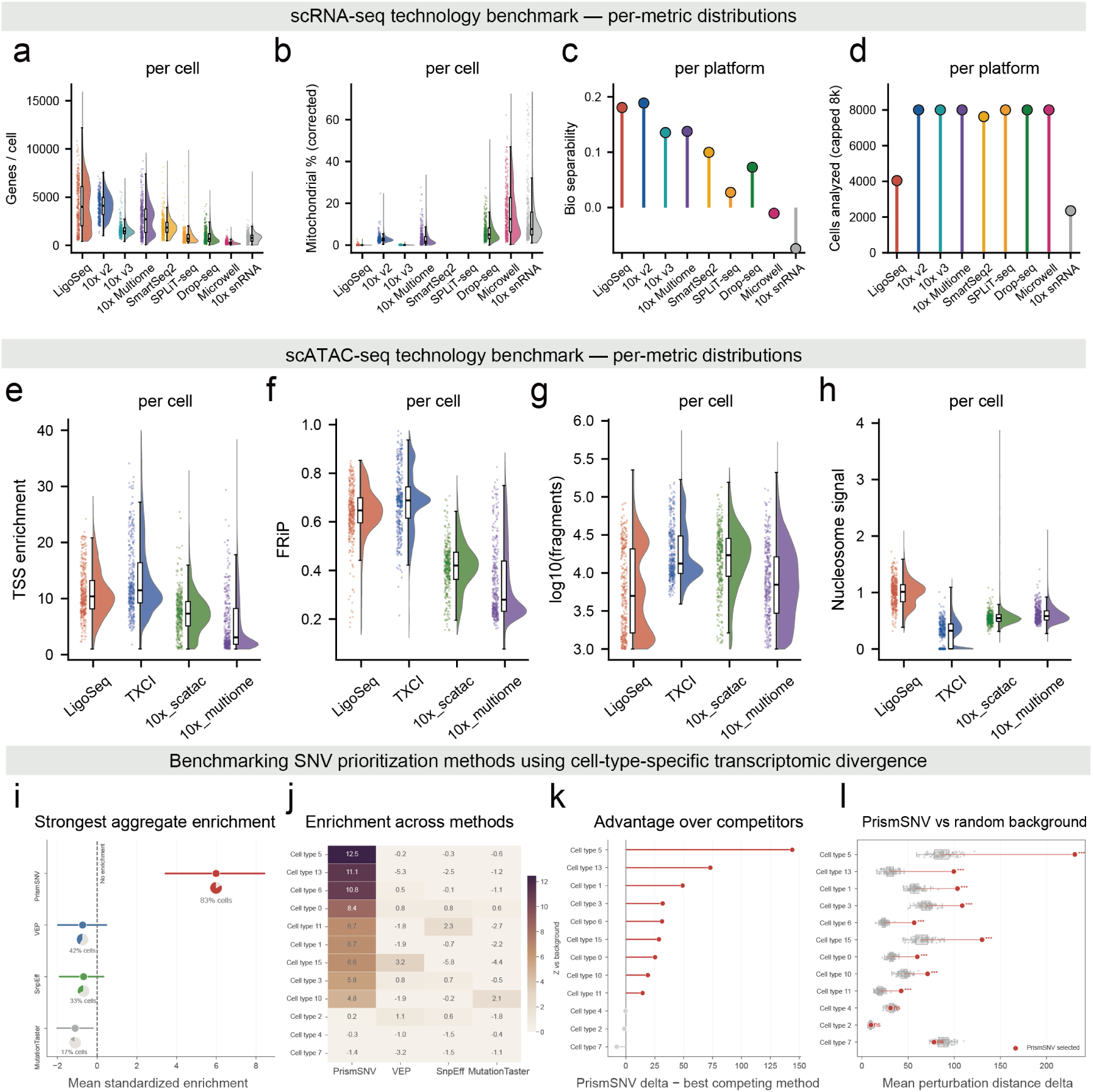
Benchmark with other technologies. **a.** Genes per cell-sensitivity (Ligo-Seq, 10x v2, 10x v3, 10x Multiome, Smart-seq2, SPLiT-seq, Drop-seq, Microwell-seq, 10x snRNA; up to 20,000 cells per platform subsampled for analysis, up to 4,000 shown; **d, e, f** used same datasets), Ligo-Seq (median 3994) matches 10x v2 (median 4106) and exceeds all other platforms. **b.** Mitochondrial%, counts in mitochondrial genes (lower is better); Ligo-Seq median 0%, versus 2.7% (10x v2) to 12.4% (Microwell-seq). Smart-seq2 and SPLiT-seq are omitted (their matrices lack annotated mouse mitochondrial genes; Methods). **c.** Bio separability-marker silhouette coefficient; highest for Ligo-Seq and 10x v2. **d.** Cells analyzed per sample-throughput context only (post-QC count, capped at 8,000); not included in the composite score. y-axes of heavy-tailed metrics are clipped at the 99th percentile for display; statistics use unclipped data. **e.** TSS enrichment benchmark (Ligo-Seq, n = 6,884; TXCI-ATAC, n = 49,758; 10x scATAC, n = 4,371; 10x Multiome, n = 48,156 QC-passing nuclei; **f, g, h** used same datasets), enrichment of fragments at transcription start sites (signal-to-noise); Ligo-Seq (median 10.4) and TXCI (median 11.5) highest, far above 10x scATAC (median 7.3) and 10x Multiome (median 3.1). **f.** FRiP-fraction of fragments in called peaks; Ligo-Seq 0.65 and TXCI 0.69 highest (10x scATAC 0.42, 10x Multiome 0.28). **g.** log10(unique fragments)-per-nucleus library depth. **h.** Nucleosome signal-mononucleosomal:nucleosome-free fragment ratio; Ligo-Seq highest (median 1.01) versus 0.55 and 0.58 for the 10x assays and 0.33 for TXCI (recomputed to account for its dual-orientation fragments file; Methods). All platforms lie far below the over-transposition exclusion threshold (>4), so higher values reflect better-resolved nucleosomal structure. **i.** Common-metric comparison of SNVs prioritized by PrismSNV, VEP, SnpEff and MutationTaster. Each method selected candidates according to its own predefined criterion: PrismSNV candidates were the top 10% ranked by perturbation score within each cell type; VEP and SnpEff candidates were classified as HIGH impact; and MutationTaster candidates were classified as Deleterious. The subsequent RNA distance-delta metric was not used for candidate selection. For every candidate set, RNA distance delta was calculated using the same procedure as the observed Euclidean distance between the mean transcriptomes of ALT- and REF-supported cells minus the mean distance obtained from randomized cell-group assignments with preserved group sizes. Mean standardized enrichment relative to random-SNV backgrounds across evaluated cell types; points denote mean Z scores, horizontal bars denote 95% bootstrap confidence intervals, and inset pie charts indicate the fraction of evaluated cell types with positive enrichment. **j.** Cell-type-level enrichment Z scores for the four prioritization methods. **k.** The difference between PrismSNV and the best-performing sequence-based method within each cell type, calculated from the observed mean RNA distance delta. **l.** Direct comparison between PrismSNV-prioritized SNVs and random-SNV backgrounds; grey points and boxplots show 100 independently sampled background sets of 100 eligible SNVs, red points show the observed mean RNA distance delta of PrismSNV-prioritized SNVs, and red segments connect the background and observed means. The same eligible SNV universe, RNA representation, distance definition and background-sampling framework were used for all methods. Asterisks denote FDR- adjusted significance: ∗ *q* < 0.05, ∗∗ *q* < 0.01, ∗∗∗ *q* < 0.001; ns, not significant.

**Supplementary Figure 2.**
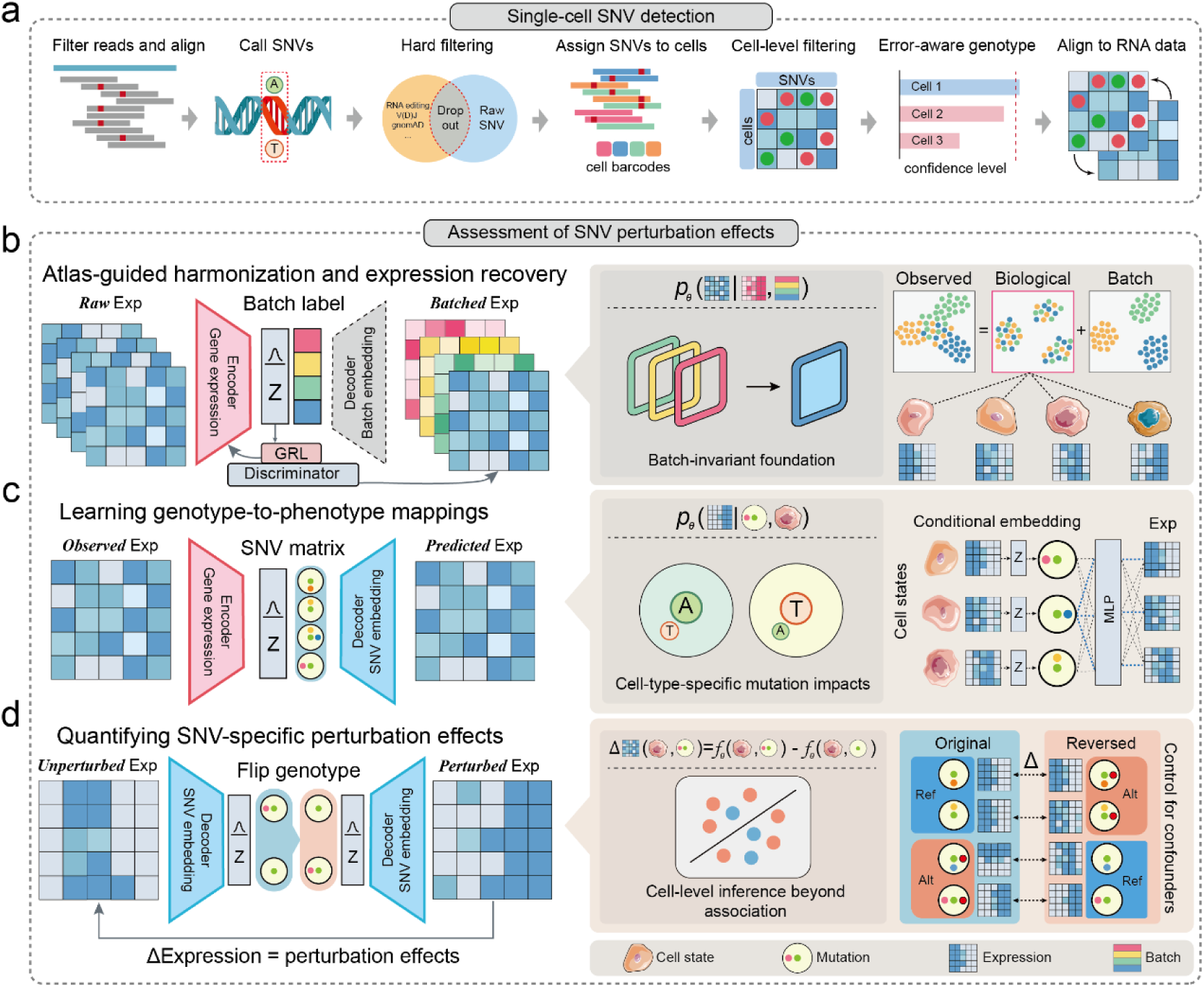
PrismSNV framework for within-cell SNV perturbation analysis. **a.** Error-aware SNV genotype extraction directly from single-cell transcriptomes, without requiring population allele frequencies, or reference annotation. Raw sequencing reads are aligned to the reference genome to identify candidate SNVs. Putative variants undergo rigorous filtering to remove RNA editing sites and technical dropouts. The filtered SNVs are subsequently assigned to individual cells via cell barcodes and integrated with matched single-cell RNA expression profiles. **b.** Batch-invariant latent space learning via adversarial variational autoencoder. A variational autoencoder maps raw gene expression data into a latent space (Z). To decouple true biological variation from technical artifacts, an adversarial discriminator equipped with a GRL is employed, ensuring that batch-associated effects are absorbed by a batch-aware decoder and eliminated from the stable latent representation. **c.** Fine-tuning phase for assessing SNV perturbation effects. Using the purified latent cellular states, the model integrates the single-cell SNV matrix to learn context-aware representations. This module captures the cell’s complete transcriptional context. **d.** By computationally toggling the allelic state of a focal mutation while holding the inferred cellular background fixed, PrismSNV estimates its perturbation effect within a matched cellular context, reducing confounding from clonal ancestry, differentiation state and microenvironmental variation.

**Supplementary Figure 3.**
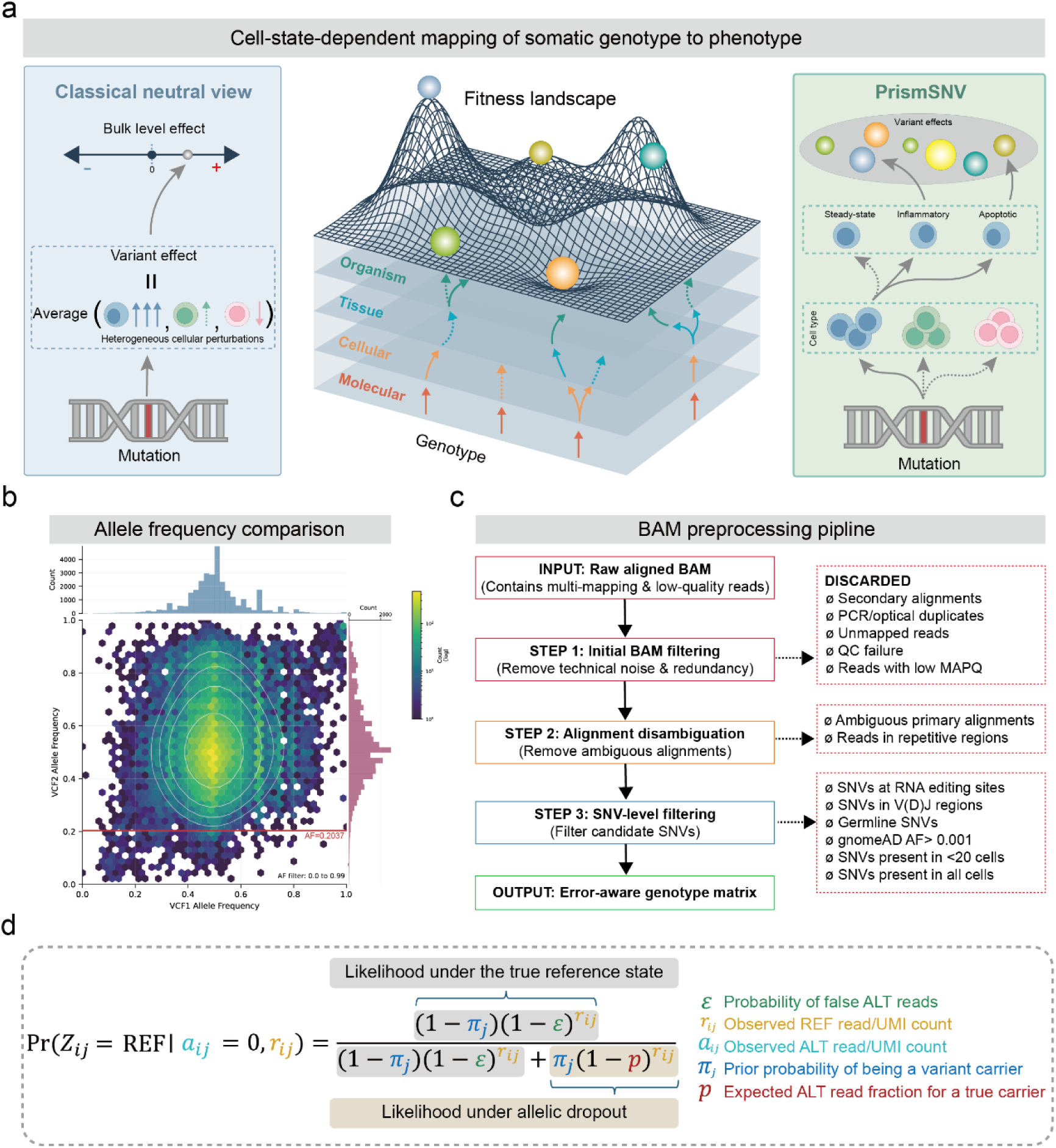
PrismSNV framework for cell-context-resolved somatic SNV genotyping and perturbation mapping. **a. Left**, the classical neutral view summarizes the effect of a somatic mutation at the population level. Although individual cells carrying the same mutation may show heterogeneous responses, these differences are collapsed into an average effect, which may obscure cell-level perturbations. **Middle**, the fitness landscape illustrates that genotype-to-fitness relationships can emerge through multiple biological layers, from molecular perturbation to cellular state, tissue context and organism-level outcomes. **Right**, PrismSNV resolves this intermediate layer by estimating variant effects within the cellular context in which the mutation is observed. The same somatic variant may produce distinct effects across cell types and cell states, remaining silent in one context but becoming perturbative in another. **b.** Comparison of allele frequencies (AFs) for matched germline SNVs detected in both whole-exome sequencing (WES, top blue histogram) and single-cell RNA sequencing (scRNA-seq, right red histogram). The central plot illustrates the density of heterozygous SNVs, overlaid with white contour lines representing the probability density. Note that homozygous variants (*AF* = 1), which account for approximately 61.2% of the sites, are excluded from this visualization to better highlight the distribution of heterozygous loci. The red horizontal line indicates the empirical filtering threshold (*AF* = 0.2037) applied to the RNA data. **c.** Schematic overview of the rigorous BAM preprocessing and SNV filtering pipeline. The workflow proceeds from raw aligned BAM files to the final error-aware genotype matrix. **d.** The Bayesian probabilistic model used to infer true cellular genotypes. The formula calculates the posterior probability that a given cell is a true reference (REF) homozygote (*Z_ijj_* = REF), given the observation that all corresponding reads or UMIs support the REF allele (*a_ijj_* = 0) and *r_ijj_* reads support the REF allele. The model explicitly accounts for single-cell technical artifacts by contrasting the likelihood of a true reference state against the likelihood of allelic dropout, incorporating key parameters such as the probability of false alternative (ALT) reads (*εε*), the prior probability of variant carriage (*π_jj_*), and the expected ALT read fraction (*p*).

**Supplementary Figure 4.**
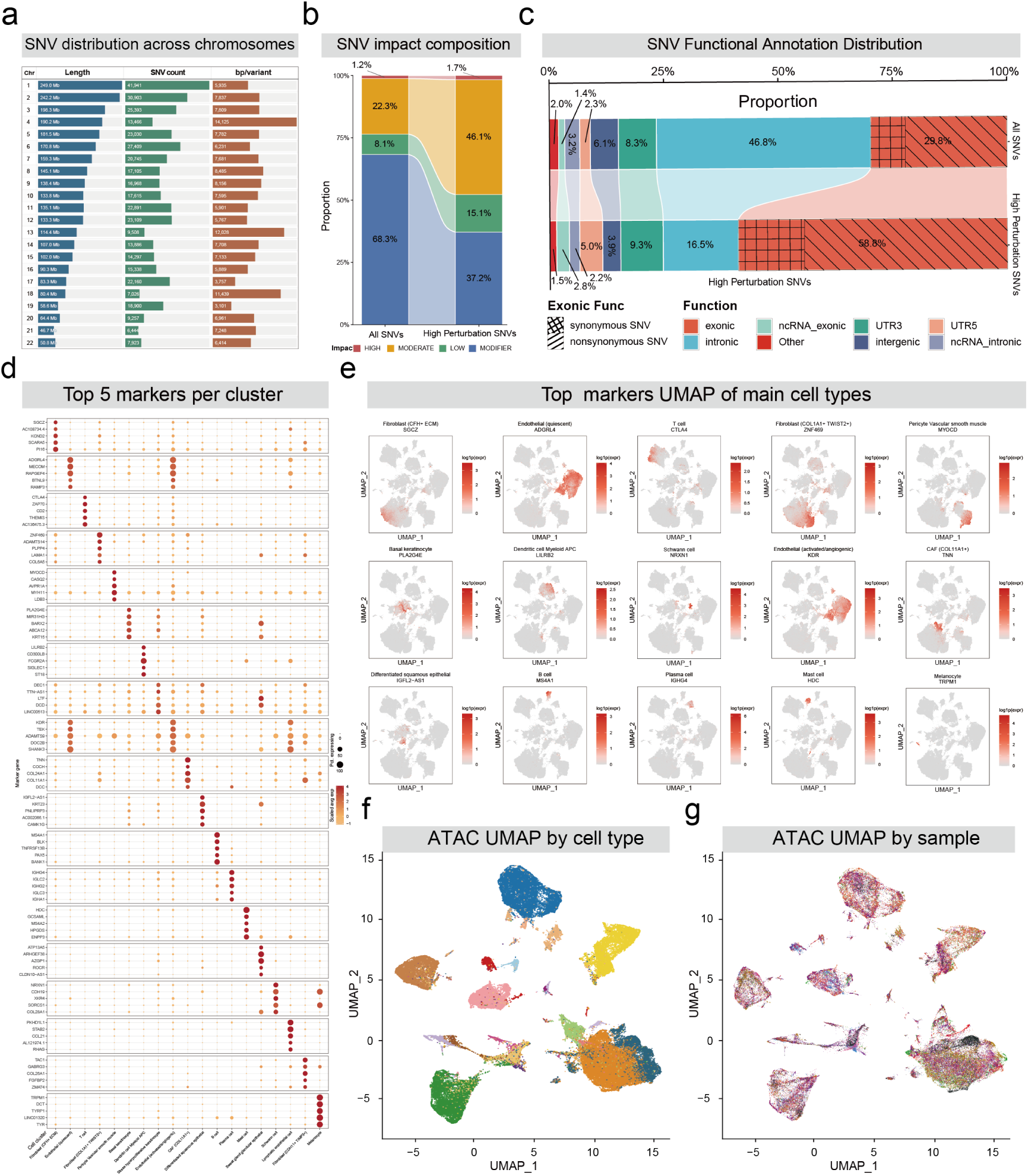
Cell-type annotation and genomic characterization of PrismSNV-detected somatic SNVs. **a.** The number and frequency of SNVs detected across different chromosomes in all full-length single-cell RNA sequencing datasets before filtering. **b.** Concordance with Variant Effect Predictor (VEP) annotations. The bar plot compares the impact composition (HIGH, MODERATE, LOW, MODIFIER) between all baseline SNVs (top) and PrismSNV-derived high-perturbation SNVs (bottom). **c.** Genomic functional distribution of all candidate SNVs (left) versus PrismSNV-identified high-perturbation SNVs (top 10% per cell type, right). **d.** Bubble plot showing the top five marker genes for each cell type in the single-cell RNA-seq dataset. **e.** UMAP visualization showing the expression patterns of representative marker genes across different cell types. **f.** UMAP visualization of the paired single-cell ATAC-seq data colored by cell type. Cell-type labels were transferred from the matched RNA-seq data. The ATAC and RNA modalities showed high concordance, with the same cell types identified in RNA-seq also forming distinct clusters in the ATAC-seq space. **g.** UMAP visualization of the paired single-cell ATAC-seq data colored by patient sample. Cell-type labels were transferred from the matched RNA-seq data. The mixed distribution of patient samples indicates effective removal of inter-sample batch effects.

**Supplementary Figure 5.**
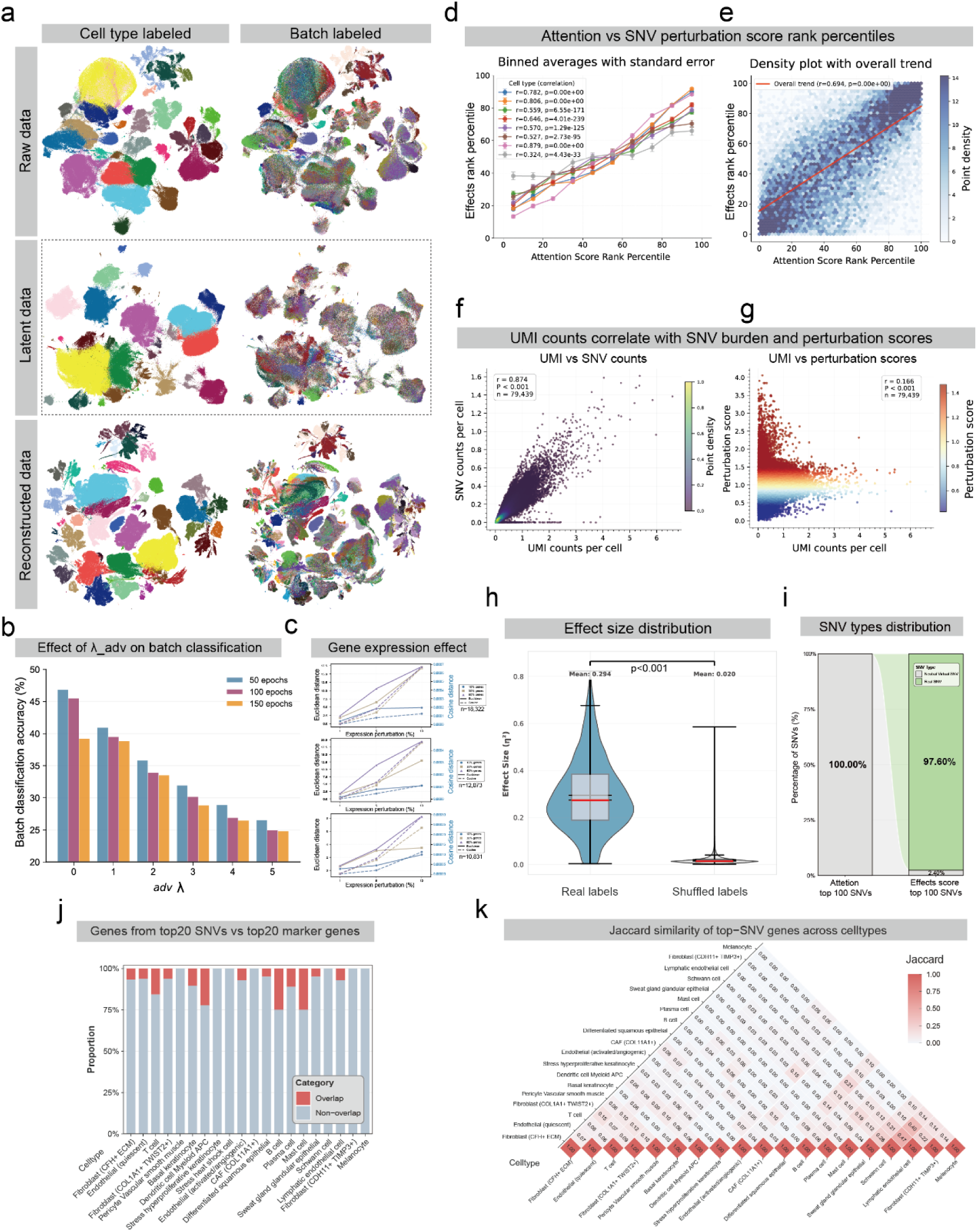
Validation of PrismSNV results. **a.** UMAP visualizations of raw expression profiles, encoder-derived latent representations, and decoder-reconstructed data, colored by cell type (left) and batch label (right). The latent space demonstrates robust batch integration compared to raw and reconstructed data. **b.** Impact of adversarial discriminator strength and training epochs on batch classification accuracy, showing effective attenuation of residual batch effects. **c.** In silico validation of perturbation metrics. Line plots confirm that Euclidean (solid lines) and cosine (dashed lines) distances accurately capture simulated transcriptomic shifts across varying perturbed gene proportions and expression magnitudes. **d, e.** Correlation between attention weights and final SNV perturbation scores, shown as binned averages per cell type (D) and a global density plot (E). PrismSNV’s attention mechanism correctly prioritizes high-impact SNVs. **f, g.** Density scatter plots ruling out sequencing depth bias. Cellular UMI counts correlate strongly with detected SNV burdens (F, r=0.874) but are independent of the derived perturbation scores (G, r=0.166). **h.** SNV-specific, sample-blocked permutation assessment of cell-type-dependent model displacement. For each candidate SNV, the observed eta-squared value was compared with 1,000 values obtained by permuting cell-type labels among covered cells within its host sample blocks. Empirical P values were calculated within SNV and adjusted across the full validation universe using the Benjamini-Hochberg procedure. The panel shows observed eta-squared values relative to their corresponding null medians or 95th percentiles. **i.** False positive evaluation using perfectly symmetric simulated variants. Synthetic SNVs monopolize the top 100 attention weights but are robustly filtered out in the final top 100 perturbation scores (accounting for 2.4%). **j.** Overlap between the top-20 PrismSNV-prioritized SNV-hosting genes and the top-20 marker genes within each cell type, quantified as the proportion of overlapping versus non-overlapping genes. **k.** Lower-triangular heatmap of Jaccard similarity between top-SNV-hosting gene sets across cell types, assessing whether prioritized genes are shared or cell-type-restricted.

**Supplementary Figure 6.**
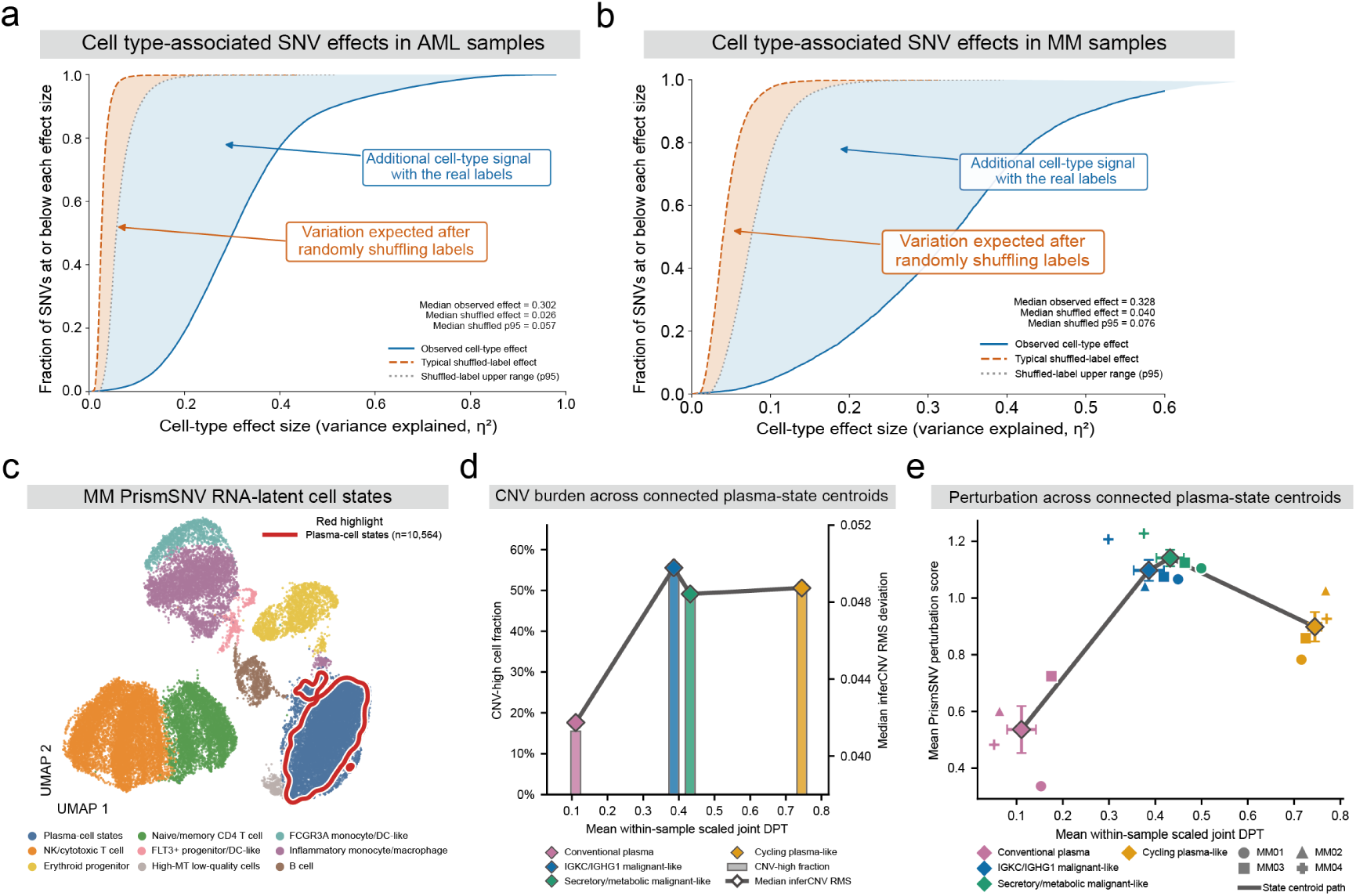
Cell-type-specific SNV perturbation effects and state-dependent displacement in AML and MM. **a.** Distribution of cell-type effect sizes of SNVs in AML samples. The horizontal axis denotes the cell-type effect size, and the vertical axis indicates the cumulative fraction of SNVs with effect sizes not exceeding the corresponding value. The blue solid line represents true cell-type labels, whereas the orange dashed and grey dotted lines represent the median and 95th percentile across 1000 within-sample label permutations, respectively (n = 37313). The light orange shaded area delineates the expected range under random label permutation, and the light blue shaded area highlights the excess signal under true labels that exceeds the permutation 95th percentile. This indicates that a subset of SNV perturbation effects exhibits cell-type-specific, heterogeneous responses rather than arising solely from stochastic label fluctuations. **b.** Distribution of cell-type effect sizes of SNVs in MM samples. The orange dashed and grey dotted lines represent the median and 95th percentile across 1000 within-sample label permutations, respectively (n=9,680). The labels in this figure are interpreted identically to those in panel **a**. **c.** UMAP projections of single-cell RNA-seq data across four MM samples, with the plasma cell compartment demarcated by red contours. **d.** CNV burden of MM plasma cell states along the within-sample normalized joint DPT. Bars indicate the proportion of CNV-high cells defined by the 95th percentile of reference cells; diamonds and the connecting line represent the median inferCNV RMS of each state (right axis); the horizontal axis denotes the mean joint DPT coordinate of each state. **e.** PrismSNV perturbation scores of MM plasma cell states along the within-sample normalized joint DPT. Different shapes denote sample-state means across individual samples. Colored diamonds represent the equal-sample-weighted centroids and SEM for the four plasma cell states, with the black line tracing the centroid trajectory. The states are sequentially arranged as conventional plasma cells, IGKC/IGHG1 malignant- like, secretory/metabolic malignant-like, and cycling plasma-like.

**Supplementary Figure 7.**
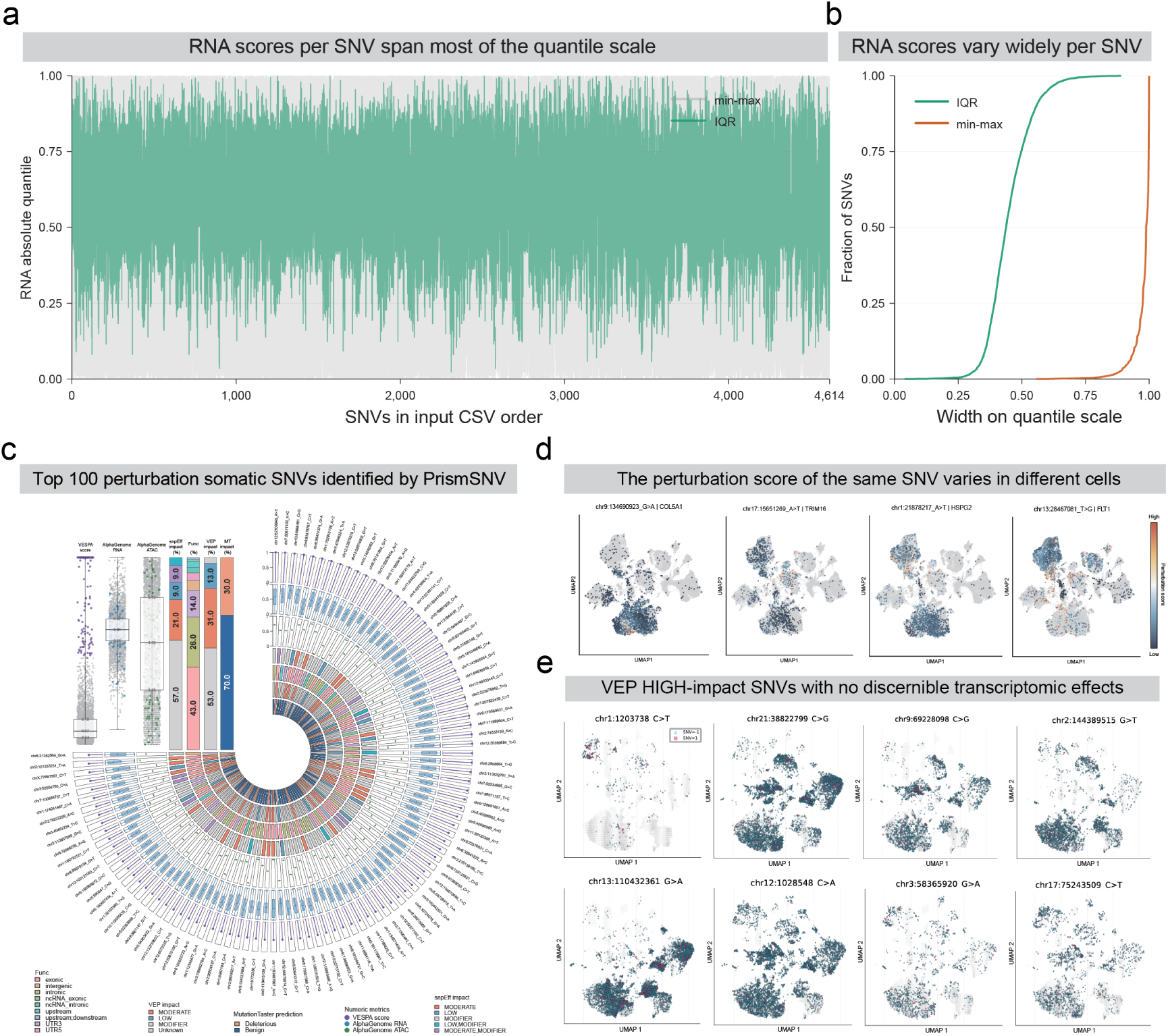
PrismSNV identifies cell-context-dependent SNV perturbations beyond sequence-based annotation. **a.** Distribution of RNA absolute quantile scores for 4,614 SNVs with available RNA summaries. For each SNV, the lighter interval denotes the minimum-to-maximum range and the darker interval denotes the interquartile range (IQR; Q1-Q3). SNVs are displayed in input-table order. **b.** Empirical cumulative distribution functions of the within-SNV IQR and minimum-to-maximum range. **c.** Prediction results from different methods for the top 100 SNVs with the largest perturbation effects identified by PrismSNV. For AlphaGenome, both RNA and ATAC queries were restricted to skin tissue contexts. The circular tracks show independent evaluation results for each SNV, whereas the upper-left panel summarizes the results across all SNVs. In the summary panel, grey points represent the remaining SNVs outside the top 100. **d.** UMAP plots for different cells show that the perturbation scores for different SNVs vary across different cells. Gray dots represent cells where the SNV was not detected. The perturbation scores are normalized from 0 to 1. Red represents high perturbation scores, and blue represents low perturbation scores. **e.** Representative SNVs classified as HIGH-impact by VEP but exhibiting no detectable perturbation in PrismSNV. Cells are color-coded by genotype: red squares indicate cells harboring the SNV, blue squares represent cells lacking the mutation, and gray squares denote cells with undetermined genotypes due to undetected signals or insufficient read coverage.

**Supplementary Figure 8.**
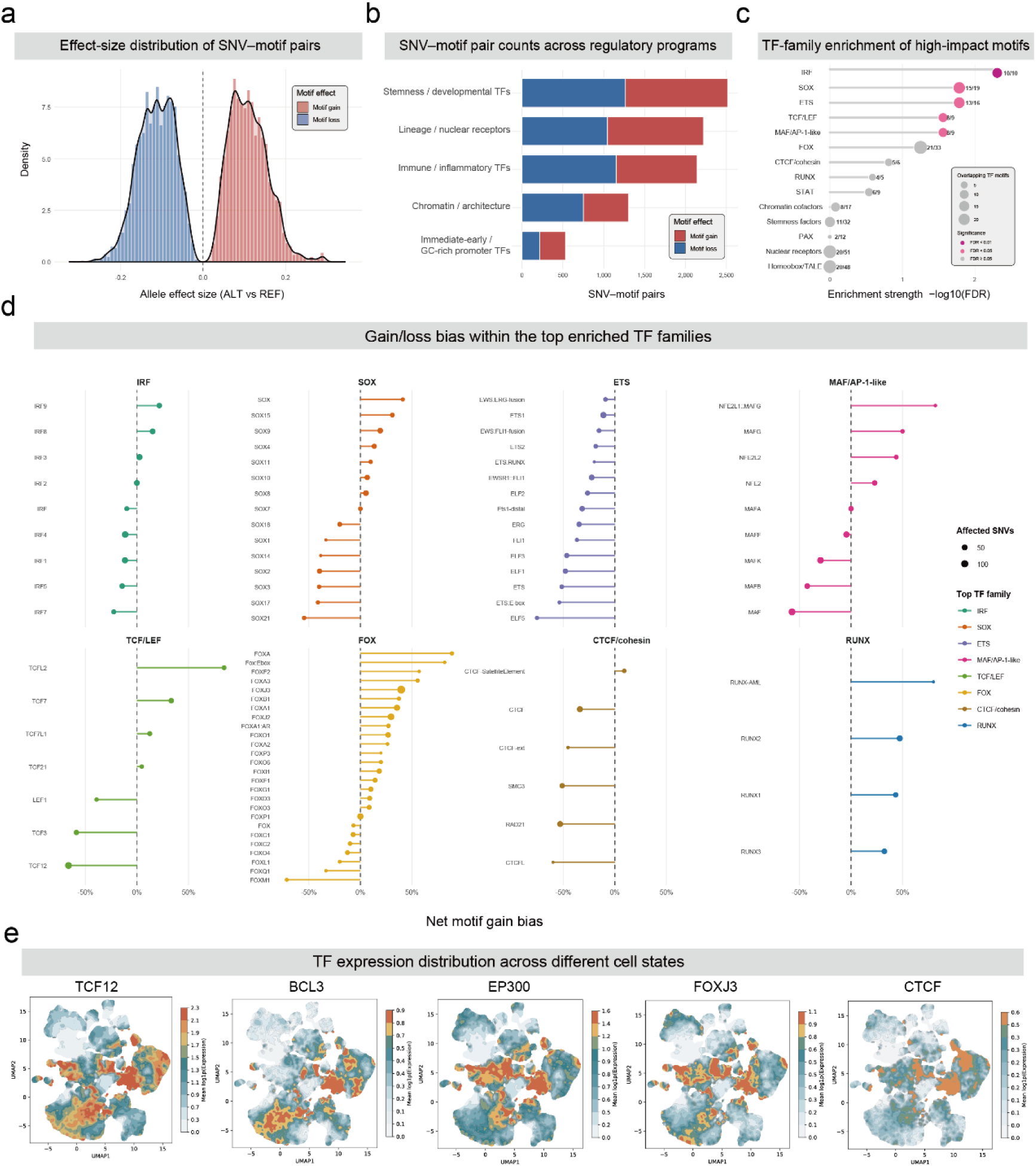
SNV-associated motif gains and losses reveal TF-family-specific regulatory perturbations. **a.** Signed motif effect-size distribution for motifbreakR-predicted SNV-motif pairs. Positive values denote stronger alternative-allele motif matching and negative values denote motif disruption; red and blue indicate motif gain and loss, respectively. **b.** Distribution of affected SNV-motif pairs across major regulatory TF programmes, stratified by predicted motif gain or loss. **c.** TF-family enrichment among high-impact motif alterations. Enrichment strength is shown as -log10(FDR); point size indicates the number of overlapping TF motifs and labels denote observed overlap over the family universe. **d.** Net gain–loss bias for TF motifs within the top enriched TF families. Net motif gain bias was calculated as (gain hits − loss hits) / total motif hits. Points to the right of zero are gain-biased and points to the left are loss-biased; point size indicates the number of affected SNVs and color denotes TF family. **e.** Differences in the expression levels of SNV-affected TF across different cell state.

**Supplementary Figure 9.**
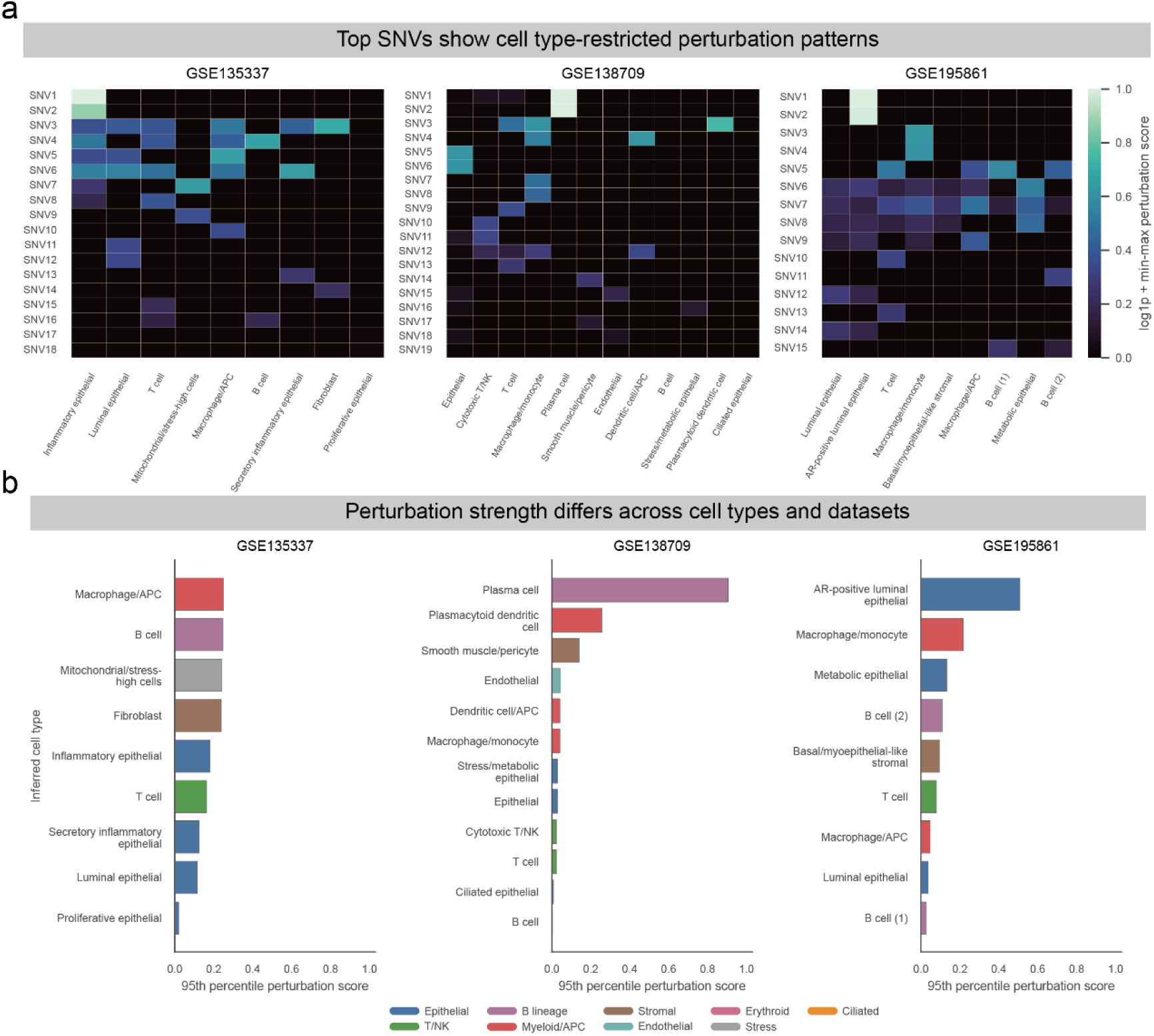
SNV perturbation effects are cell-type specific across independent datasets. **a.** Heat maps showing the perturbation scores of top-ranked SNVs across inferred cell types in GSE135337, GSE138709 and GSE195861. Rows denote individual SNVs and columns denote inferred cell types. Color intensity represents the log1p-transformed perturbation score after min-max normalization, with higher values indicating stronger predicted transcriptomic perturbation. Top SNVs displayed sparse and cell type-restricted activity patterns rather than uniform effects across all cell populations. **b.** Cell type-level perturbation burden summarized by the 95th percentile of per-cell perturbation scores in each dataset. Bars are grouped by dataset and colored according to broad cellular lineages. The strongest perturbation signals were concentrated in specific epithelial, immune and stromal compartments, indicating that SNV-associated effects vary substantially across cell types and datasets.

